# Negative HIV serology in perinatally infected children after early treatment and viral suppression: an exploratory analysis of immune correlates

**DOI:** 10.64898/2026.05.29.728638

**Authors:** Pierre Frange, Jerôme Le Chenadec, Daniel Scott-Algara, Caroline Charre, Thomas Montange, Elise Gardiennet, Ingrid Fert, Adeline Mélard, Damien Batalie, Stéphane Blanche, Catherine Dollfus, Marie-Dominique Tabone, Albert Faye, Josiane Warsawski, Véronique Avettand-Fenoël, Florence Buseyne, the ANRS-EP59-Study Group

## Abstract

**Background:** Undetectable HIV-specific antibodies in early-treated children with confirmed infection correlate with low viral reservoir and may identify those eligible for future HIV remission strategies. The neonatal immune system’s unique characteristics, combined with impairments resulting from exposure to maternal HIV and antiretroviral treatment (ART), may affect antibody responses to HIV. Yet immune competence remains understudied in the context of negative HIV serology. The ANRS-EP59-CLEAC study included 76 children and adolescents with HIV. We measured plasma HIV antibodies by enzyme immunoassay, other analytes by ELISA or multiplex assays, and blood cell phenotypes and functions by flow cytometry. We used Fisher’s and Mann-Whitney’s tests and logistic regression to analyze variables associated with negative HIV serology. Nine participants tested negative for HIV-specific antibodies, eight children and one adolescent. Negative HIV serology occurred exclusively in participants who had initiated ART early and had HIV RNA < 50 copies/mL at evaluation. Among 17 early-treated children with sustained viral suppression, only 7 had negative HIV serology. In this subgroup, negative HIV serology associated with higher nadir CD4 counts, lower plasma IgM levels, higher frequencies of circulating follicular CD8 T lymphocytes, and higher expression of the costimulatory molecule CD86 on myeloid dendritic cells. We found no evidence of B or T lymphocyte deficits associated with negative HIV serology. Low antigenic exposure was necessary but insufficient to explain negative HIV serology. Beyond its association with low HIV reservoir, negative HIV serology correlated with less severe prior CD4 T-lymphocyte depletion and higher frequencies of follicular CD8 T lymphocytes.

**Summary:** In HIV-infected infants, starting antiretroviral therapy (ART) very early dramatically improves health outcomes. An important phenomenon observed in some of these children is that, despite being HIV-infected, they show no detectable antibodies against the virus — a profile referred to as negative HIV serology. This feature could help identify patients most likely to benefit from future strategies aimed at achieving long-term virus control without treatment. To better understand this phenomenon, we studied HIV-infected children and adolescents enrolled in the ANRS-EP59-CLEAC trial, which compares the effects of ART initiated early (before 6 months of age) versus late (after 24 months). We showed that negative HIV serology is associated with early treatment, but also with a better-preserved immune system: less depletion of CD4 T cells, which are critical immune cells, and a higher abundance of specific T lymphocytes with potent antiviral activity. Importantly, we found no evidence of defects in the mechanisms responsible for antibody production. These findings suggest that negative HIV serology reflects a favorable immunological profile and could serve as a useful marker to select children as candidates for HIV remission trials.

## Introduction

Maternal antiretroviral treatment (ART) has dramatically reduced mother-to-child HIV transmission, yet over 120,000 children infections occur annually [1]. Early ART in infected infants reduces morbidity and mortality while limiting reservoir seeding and immune alterations [2]. Despite these benefits, lifelong adherence remains challenging, making treatment-free remission an urgent research priority [2]. Achieving this goal requires understanding immunological and virological characteristics of early-treated children.

An important phenomenon in early-treated children is negative HIV serology: after clearance of *in utero* transmitted maternal antibodies, HIV-specific antibodies are undetectable despite confirmed infection [3–7]. This discordance between serology and infection status, as given by current or previous positive PCR results occurs in up to half of early-treated children [5–8], which poses challenges for clinical management, and confuses perinatally infected children, youths, and their caregivers [6,8]. This confusion may reduce ART adherence [6]. Yet, negative HIV serology represents an opportunity because it associates with low HIV reservoirs and may serve as a useful and simple clinical marker to identify patients eligible for future HIV remission strategies [2,7].

Most reports on negative HIV serology have not addressed immune competence, despite its importance during perinatal HIV exposure. This importance stems from fundamental differences between the neonatal and adult immune systems: neonatal immunity shows attenuated responses against pathogens to avoid immunopathological consequences [9]. In addition to these developmental characteristics, exposure to maternal HIV and/or antiretroviral drugs impairs immune system development [10–12]. In HIV-infected neonates, decreased B-cell numbers and abnormal B-cell activation at birth may affect their antibody response to HIV [13]. Finally, while many HIV-infected children benefit from ART, a significant proportion experience treatment interruption during childhood or adolescence; in these cases, absent HIV-specific immune responses may lead to greater viral rebound [4].

The ANRS-EP59-CLEAC study investigates the long-term impact of age at ART initiation (< 6 months vs. ≥ 24 months of age) on immunological and virological characteristics of HIV-1-infected children and adolescents living in France [14,15]. Here, we focus on negative HIV serology, determined by a fourth-generation enzyme immunoassay (EIA). Although negative HIV serology strongly associated with early treatment and viral suppression overall, only 41% (7/17) of early-treated, virally suppressed children were seronegative at ages 5-12 years. This suggests that factors other than viral exposure determine serological responses. We present an exploratory analysis of these factors.

## Methods

### Participants and biological samples

We previously described the 76 children and adolescents included in the ANRS-EP59-CLEAC study between June 20^th^ 2016 and January 30^th^ 2019 [14,15]; their characteristics are presented in S2-1 Table. We conducted the study in accordance with the Declaration of Helsinki, and the “Comité de protection des personnes Île-de-France V” approved the protocol. The ClinicalTrials.gov identifier is NCT02674867. Participants agreed to participate if they were old enough to provide informed assent, and at least one parent provided written informed consent. We included participants who met the following criteria: (1) acquired HIV-1 through vertical transmission, (2) were aged 5 to 17 years at inclusion, (3) initiated ART for therapeutic purposes before six months of age (Early treatment group) or after two years of age (Late treatment group), and (4) achieved initial virological success (HIV-1 RNA < 400 copies/mL within 24 months of treatment initiation), regardless of subsequent viremia. We collected blood in EDTA tubes between 2016 and 2019 for biological evaluations.

### Virological assays

We performed a fourth-generation HIV EIA (VIDAS® HIV DUO ULTRA, bioMérieux) on plasma samples. Samples were considered positive when values exceeded 0.25 arbitrary units, according to the manufacturer’s instructions. We quantified total cell-associated HIV DNA using ultrasensitive real-time PCR (Biocentric) in whole blood [14,16]. We extracted HIV DNA from 1 mL of whole blood and tested it in two to four PCRs. We reported results as either actual HIV DNA copy numbers/10⁶ PBMCs or, when cellular HIV DNA fell below the detection threshold, as an arbitrary value corresponding to half the threshold value. We calculated thresholds for each assay, which varied according to available cell numbers and ranged from 11 to 24 copies/10⁶ PBMCs. We quantified integrated HIV DNA using ultrasensitive Alu-PCR as previously described [17], with thresholds ranging from 10 to 38 copies/10⁶ PBMCs. We used the Liaison XL CMV IgG (Diasorin) chemiluminescence immunoassay to quantify cytomegalovirus (CMV) antibodies.

### Immunological investigations

We quantified soluble analytes in plasma samples using commercial ELISA, Luminex, or SIMOA kits, according to the manufacturers’ instructions. For Luminex kits, we prepared and analyzed samples with DropArray™ 96 plates (Curiox) and recorded fluorescence on a Bioplex 200 (Biorad). For SIMOA kits, we recorded fluorescence on an HD-X analyzer (Quanterix). S1-1 Table describes the reference, quantification threshold, and number of samples with analyte levels below the quantification threshold or whose values were extrapolated because the signal was out of the quantification range.

We quantified lymphocyte and myeloid cell subsets in fresh blood by flow cytometry. We stained whole blood with antibody mixtures in 5 mL tubes and incubated cells for 30 min at room temperature (RT) in the dark. We then added 3 mL of lysis solution (eBioscience, 00-5333-57), incubated cells for 15 min at RT, washed them once in PBS, and immediately analyzed them with a Gallios flow cytometer (Beckman-Coulter). Supplementary material describes staining conditions, gating strategies, and cell subset definitions.

We performed second-field (4-digit)–resolution human leukocyte antigen-B genotyping using Luminex reverse PCR sequence-specific oligonucleotides (Canoga Park, CA).

We used frozen PBMCs to detect memory B-lymphocyte expansion and anti-HIV IgG production. We stimulated them with R848 (500 ng/mL, Miltenyi 130-109-376) and IL-2 (100 U/mL, Miltenyi 130-097-744), supplementing half the wells with IL-6 (10 ng/mL, Miltenyi 130-095-352). On day 7 post-stimulation, we harvested cells from three wells to quantify B lymphocytes and CD27^hi^CD38^hi^CD21^lo^CD19^+^ plasmablasts by flow cytometry (supplementary material). We maintained two wells in culture and collected supernatants on day 12 post-stimulation. We quantified total and HIV-specific IgG concentrations in the culture supernatants by ELISA; for the latter we used recombinant HIV proteins as antigens (1 µg/mL HIV-1 IIIB p24, #12028; 0.5 µg/mL HIV-1 UG037 gp140, #12063; 0.5 µg/mL HIV-1 IIIB gp120, #12027; NIH AIDS Reagent Program).

We performed IFN-γ-based ELISpot on samples from 36 participants with sufficient fresh PBMCs, as previously described (supplemental materials and [15]). We stimulated cells with HIV-1 potential T-lymphocyte epitope (PTE) Gag and Nef peptide pools (NIH AIDS Reagent Program, #12437 and #12822) and HCMV pp65 peptide pool (#11549) at a final concentration of 1 µg/mL per peptide. We used cells incubated in medium and peptide diluent (0.2% DMSO, Sigma-Aldrich D4540) as negative controls and cells stimulated with phorbol myristate acetate (25 ng/mL, Sigma-Aldrich Chimie, P8139) and ionomycin (100 ng/mL, Sigma-Aldrich Chimie, I3909) as positive controls. We determined the number of IFN-γ-producing cells using an S6 Ultimate Image UV analyzer (CTL Europe, Bonn, Germany). We required a minimum of 10 spots/well to calculate IFN-γ-producing cell frequencies, corresponding to a detection threshold of 50 IFN-γ-producing cells/10⁶ PBMCs.

We quantified intracellular cytokine production in frozen PBMCs stimulated with the same peptide pools used for the ELISpot assay. We used cells stimulated with Dynabeads® Human T-Activator CD3/CD28 (5 µL/well, Gibco, 11132D) as positive controls. We added Brefeldin A (4 µg/mL, BioLegend, #460201) to block cytokine secretion and incubated cells overnight at 37°C in 5% CO₂. We then stained cells with antibodies targeting CD3, CD4, CD8, and CD40L and a viability dye for 15 min at RT. After fixation and permeabilization using the Fix & Perm kit (Invitrogen, GAS003), we stained cells with antibodies against IL-2, TNF-α, MIP-1β, and IFN-γ for 15 min at RT. We collected data on a CytoFlex cytometer (Beckman Coulter) and analyzed them with the Kaluza software (Beckman Coulter). The gating strategy sequentially defined lymphocytes, then CD3^+^ T lymphocytes, and finally CD4 and CD8 T lymphocytes expressing each cytokine (see supplemental materials). For CD4 T lymphocytes, we additionally assessed CD40L co-expression with each cytokine to identify antigen-responsive cells. We set the positivity threshold at 0.1% cytokine-positive cells, based on background responses in uninfected donors.

### Statistics

To account for differences in age and cumulative ART exposure among participants, we normalized variables quantifying the duration of exposure to HIV replication/suppression for age (a proxy for lifetime exposure) or for time since first combined ART (ART1) initiation. Cumulative viremia is the time-updated area under the curve of plasma HIV RNA levels [18]. Normalized cumulative viremia is the cumulative viremia divided by the relevant period of time (since first ART initiation or over the last five years before sampling), expressed as log_10_ HIV RNA copies/mL. Clinical sites systematically tested HIV RNA after ART initiation for all children. To quantify lifetime exposure to HIV replication, and to account for missing plasma HIV RNA measurements between birth and first ART initiation in some participants, we assumed that all children had detectable HIV replication from the time of infection until first ART-induced viral suppression. HIV RNA assay thresholds used by various laboratories changed during participants’ follow-up, and we diluted some plasma samples when blood volumes were insufficient. Therefore, we used the 50 copies/mL cutoff for HIV RNA measurements obtained over the last five years of follow-up and the 400 copies/mL cutoff for lifetime HIV RNA measurements. Before ART1 initiation, CD4 T-cell measurements were missing for 23% of participants; therefore, we assessed past immunological status using the period after ART1. We compared quantitative and qualitative variables between groups using the Mann-Whitney and Fisher’s exact tests, respectively. We used the Spearman test to assess correlations between quantitative variables. We used Firth’s logistic regression with added covariate to assess associations between negative HIV serology and independent variables because of the small sample size [19]. A *P* value of <.05 was used to define statistical significance. We performed analyses with Stata 16.0 and the logistf package from R (https://cran.r-project.org/web/packages/logistf/index.html).

## Results

### Low exposure to antigenic stimulation was necessary but insufficient to explain negative HIV serology

Among the 76 ANRS-EP59-CLEAC study participants, we identified nine with negative HIV serology (seronegative, SN, 12%) and 67 with positive HIV serology (seropositive, SP, 88%) using a fourth-generation HIV EIA. Eight SN were children (ages 5-12 years) and one was an adolescent (age 15 years) (17% vs 3%, *P* = 0.08). All SN belonged to the early treatment group (25% vs 0%, *P* = 0.0007).

All SN had HIV RNA < 50 copies/mL at the time of the study. We quantified total and integrated cell-associated HIV DNA [20,21]. SN had significantly lower total HIV DNA levels than SP (medians: 1.90 vs 2.82 log_10_ copies/10⁶ PBMCs, *P* = 0.0003, Figure 1A). None of the SN had detectable integrated HIV DNA (SN vs SP: 0% vs 24%, *P* = 0.19). SN had a shorter duration of exposure to HIV replication than SP, as reflected by a younger age at viral suppression <4 00 copies/mL (Age VS, 137 vs 1196 days, *P* = 0.002, Figure 1B) and a shorter lifetime exposure to HIV RNA > 400 copies/mL (11 vs 60 months, *P* = 0.0002, Figure 1C). To quantify viremia over time, we calculated time-normalized cumulative viremia (cumulative viremia divided by the duration of the analyzed time window). Normalized cumulative viremia did not differ significantly between groups over the period between ART1 initiation and the study (0.04 vs 0.14 log_10_ HIV RNA copies/mL, *P* = 0.22, Figure 1D) but SN had lower normalized cumulative viremia over the last five years before the study than SP (0 vs 0.009 log_10_ HIV RNA copies/mL, *P* = 0.003, Figure 1E).

**Figure 1.**
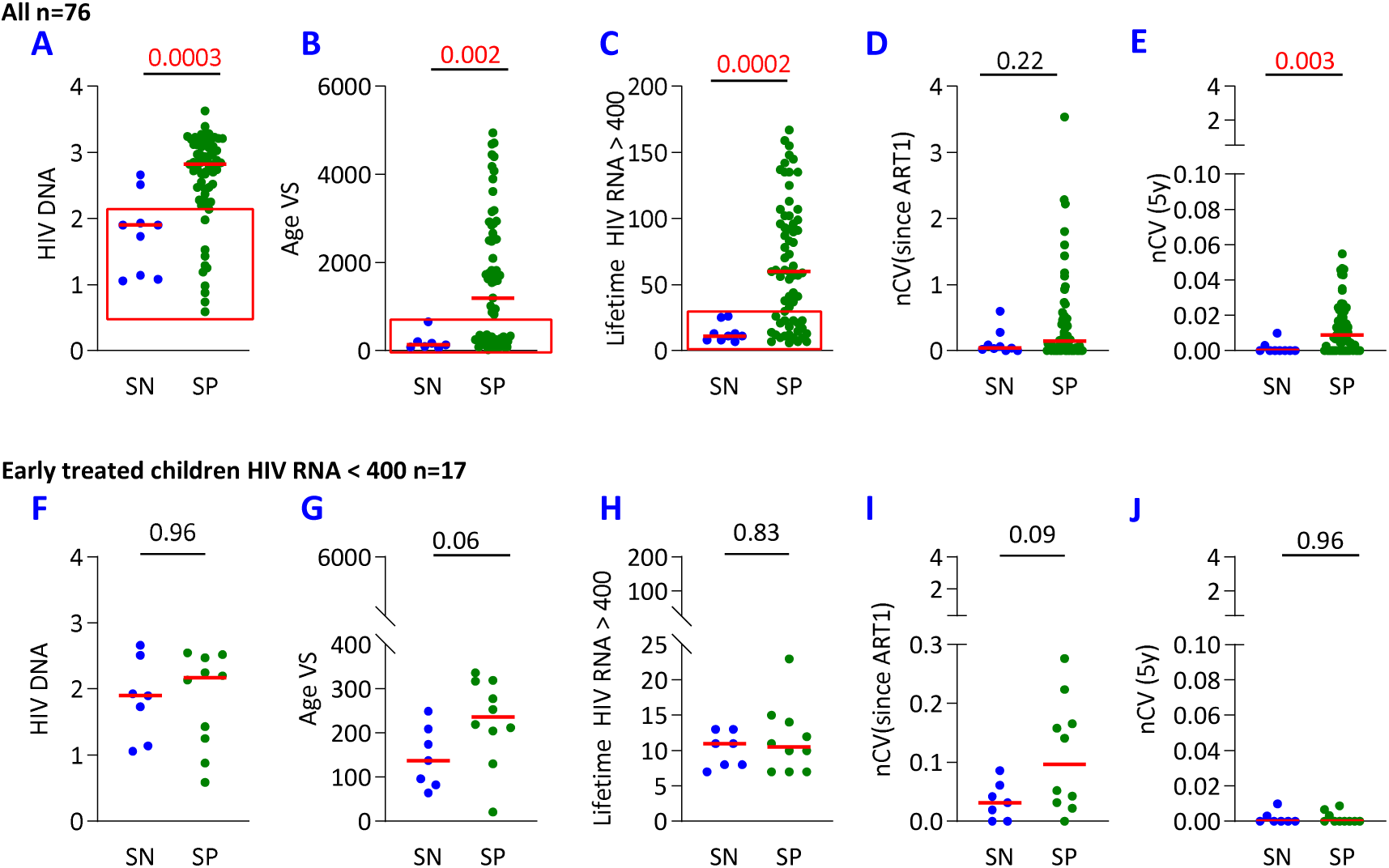
Negative HIV serology and exposure to HIV replication. We determined HIV serology in all 76 participants using a fourth-generation HIV enzyme immunoassay (EIA). We compared variables characterizing exposure to HIV replication between SN and SP participants, either across the full cohort (A to E) or in the 17 early-treated children with sustained HIV suppression (F to J). Panels show: A, F: total HIV DNA in PBMCs (log_10_ copies/10^6^ PBMCs); B, G: age at viral suppression defined as age at first HIV RNA < 400 copies/mL (days); C, H: lifetime duration of HIV RNA > 400 copies/mL (months); D, I: normalized cumulative viremia after ART1 initiation (log_10_ HIV RNA copies/mL); E, J: normalized cumulative viremia over the last five years (log_10_ HIV RNA copies/mL). Red bars indicate medians and *P* values above graphs correspond to Mann-Whitney tests. Red boxes on panels A to C highlight the overlapping distribution of exposure to HIV replication between SN and SP groups.

However, low exposure to HIV replication did not necessarily associate with negative HIV serology, as virological parameters overlapped between SN and SP. Indeed, among 17 participants with total HIV DNA < 2 log₁₀ copies/10⁶ PBMCs, seven were SN and ten were SP (highlighted with a red box in Figure 1A). Thirty participants had a lifetime duration with HIV RNA > 400 copies/mL of < 30 months, including nine SN and 21 SP (Figure 1B). Similarly, 35 participants reached HIV RNA < 400 copies/mL during their first year of life, comprising eight SN and 25 SP (Figure 1C). These observations support that although low antigenic stimulation resulting from limited HIV replication was necessary, it was insufficient to account for negative HIV serology in perinatally infected participants receiving early ART.

To identify factors associated with negative HIV serology other than exposure to viral antigens, we focused subsequent investigations on the 17 early-treated children with HIV RNA < 50 copies/ml at the time of the study who never experienced a viral rebound > 400 copies/mL after ART1 initiation. Seven were SN and 10 were SP. We found no significant differences between SN and SP in three biological measures of viral exposure: total HIV DNA (log₁₀ copies/10⁶ PBMCs: 1.9 vs. 2.2, *P* = 0.96, Figure 1F), integrated HIV DNA detection (0% vs 40%, *P* = 0.10), and plasma HIV-1 p24 detection (14% vs 30%, *P* = 0.60, Table 1). SN tended to be younger than SP at ART1 initiation (14 vs. 97 days, *P* = 0.06) and at viral suppression (137 vs 236 days, *P* = 0.06, Figure 1G). We found no significant differences in past exposure to HIV replication between SN and SP participants, as measured by lifetime duration with HIV RNA > 400 copies/ml, normalized cumulative viremia since ART1 initiation, or over the last five years (Figure 1H-J). These analyses support that negative HIV serology did not associate with lifetime or current measures of HIV exposure in early-treated children with sustained viremia suppression. Thus, additional factors associated with negative HIV serology should be identified in this population.

**Table 1.**
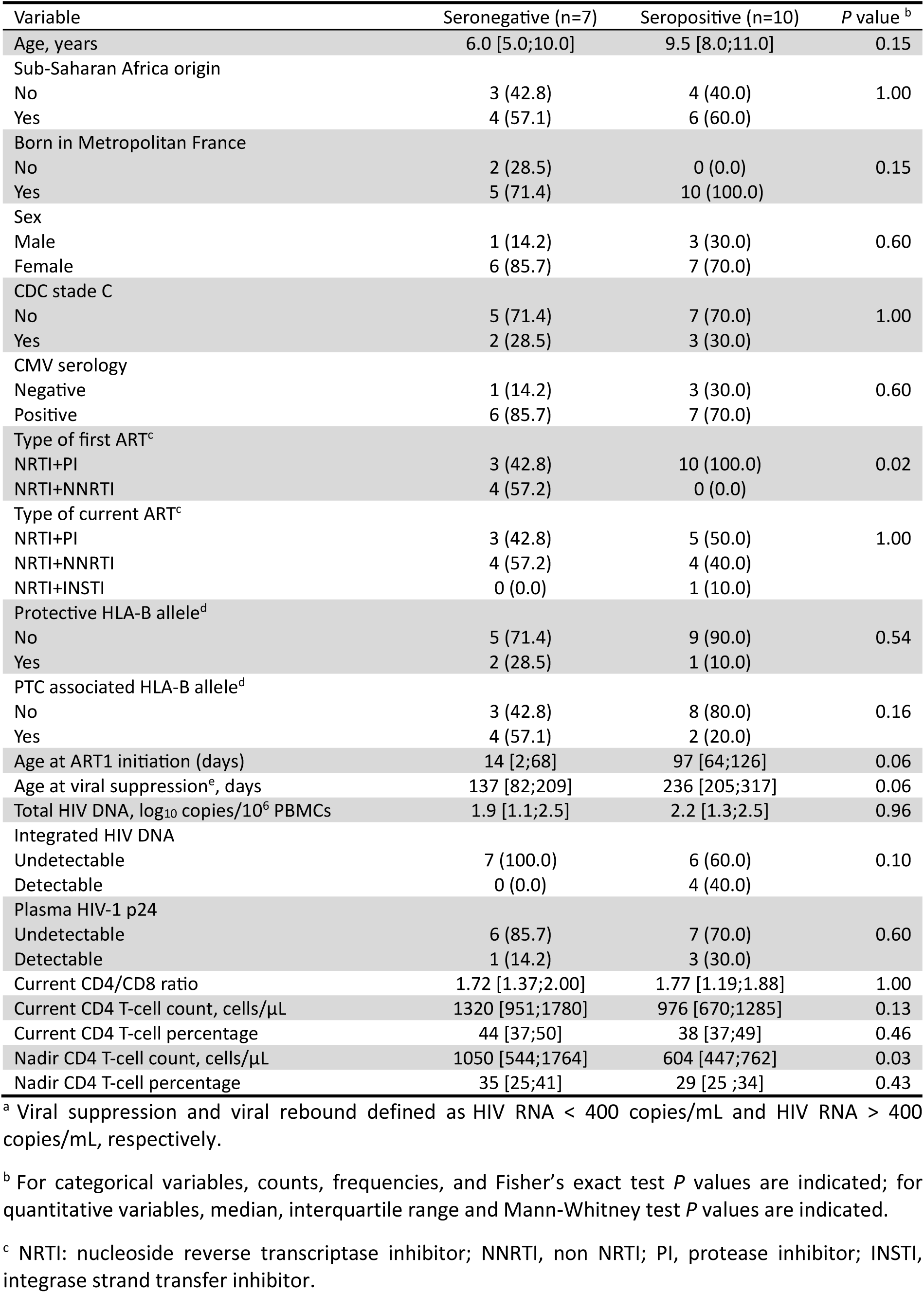
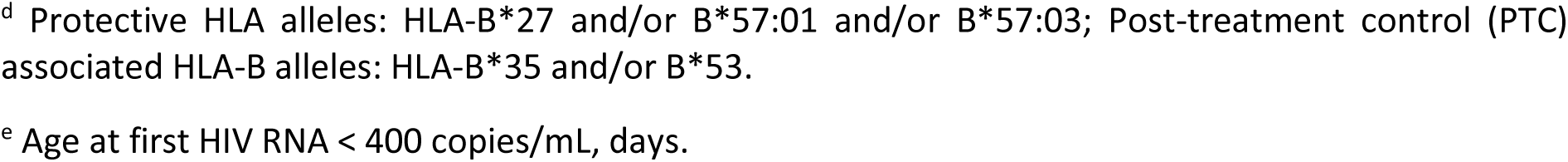
Characteristics of the 17 children who initiated ART before 6 months of age, reached viral suppression and never experienced a viral rebound thereafter ^a^.

| Variable | Seronegative (n=7) | Seropositive (n=10) | P value <sup>b</sup> |
| --- | --- | --- | --- |
| Age, years | 6.0 [5.0;10.0] | 9.5 [8.0;11.0] | 0.15 |
| Sub-Saharan Africa origin |  |  |  |
| No | 3 (42.8) | 4 (40.0) | 1.00 |
| Yes | 4 (57.1) | 6 (60.0) |  |
| Born in Metropolitan France |  |  |  |
| No | 2 (28.5) | 0 (0.0) | 0.15 |
| Yes | 5 (71.4) | 10 (100.0) |  |
| Sex |  |  |  |
| Male | 1 (14.2) | 3 (30.0) | 0.60 |
| Female | 6 (85.7) | 7 (70.0) |  |
| CDC stade C |  |  |  |
| No | 5 (71.4) | 7 (70.0) | 1.00 |
| Yes | 2 (28.5) | 3 (30.0) |  |
| CMV serology |  |  |  |
| Negative | 1 (14.2) | 3 (30.0) | 0.60 |
| Positive | 6 (85.7) | 7 (70.0) |  |
| Type of first ART<sup>c</sup> |  |  |  |
| NRTI+PI | 3 (42.8) | 10 (100.0) | 0.02 |
| NRTI+NNRTI | 4 (57.2) | 0 (0.0) |  |
| Type of current ART<sup>c</sup> |  |  |  |
| NRTI+PI | 3 (42.8) | 5 (50.0) | 1.00 |
| NRTI+NNRTI | 4 (57.2) | 4 (40.0) |  |
| NRTI+INSTI | 0 (0.0) | 1 (10.0) |  |
| Protective HLA-B allele<sup>d</sup> |  |  |  |
| No | 5 (71.4) | 9 (90.0) | 0.54 |
| Yes | 2 (28.5) | 1 (10.0) |  |
| PTC associated HLA-B allele<sup>d</sup> |  |  |  |
| No | 3 (42.8) | 8 (80.0) | 0.16 |
| Yes | 4 (57.1) | 2 (20.0) |  |
| Age at ART1 initiation (days) | 14 [2;68] | 97 [64;126] | 0.06 |
| Age at viral suppression<sup>e</sup>, days | 137 [82;209] | 236 [205;317] | 0.06 |
| Total HIV DNA, log<sub>10</sub> copies/10<sup>6</sup> PBMCs | 1.9 [1.1;2.5] | 2.2 [1.3;2.5] | 0.96 |
| Integrated HIV DNA |  |  |  |
| Undetectable | 7 (100.0) | 6 (60.0) | 0.10 |
| Detectable | 0 (0.0) | 4 (40.0) |  |
| Plasma HIV-1 p24 |  |  |  |
| Undetectable | 6 (85.7) | 7 (70.0) | 0.60 |
| Detectable | 1 (14.2) | 3 (30.0) |  |
| Current CD4/CD8 ratio | 1.72 [1.37;2.00] | 1.77 [1.19;1.88] | 1.00 |
| Current CD4 T-cell count, cells/μL | 1320 [951;1780] | 976 [670;1285] | 0.13 |
| Current CD4 T-cell percentage | 44 [37;50] | 38 [37;49] | 0.46 |
| Nadir CD4 T-cell count, cells/μL | 1050 [544;1764] | 604 [447;762] | 0.03 |
| Nadir CD4 T-cell percentage | 35 [25;41] | 29 [25 ;34] | 0.43 |

<sup>a</sup> Viral suppression and viral rebound defined as HIV RNA < 400 copies/mL and HIV RNA > 400 copies/mL, respectively.
<sup>b</sup> For categorical variables, counts, frequencies, and Fisher's exact test *P* values are indicated; for quantitative variables, median, interquartile range and Mann-Whitney test *P* values are indicated.
<sup>c</sup> NRTI: nucleoside reverse transcriptase inhibitor; NNRTI, non NRTI; PI, protease inhibitor; INSTI, integrase strand transfer inhibitor.
<sup>d</sup> Protective HLA alleles: HLA-B\*27 and/or B\*57:01 and/or B\*57:03; Post-treatment control (PTC) associated HLA-B alleles: HLA-B\*35 and/or B\*53.
<sup>e</sup> Age at first HIV RNA < 400 copies/mL, days.

### Negative HIV serology associated with higher nadir CD4 T-cell counts

We searched for demographic, genetic, and HIV infection history factors that might associate with negative HIV serology. We found no differences between SN and SP in ethnic origin, place of birth, sex ratio, age at the time of the study, previous CDC stage C events, CMV serology, or distribution of HLA-B alleles linked with disease progression in adults (Table 1). Notably, SN more frequently expressed HLA-B alleles associated with post-treatment control, but this difference did not reach significance (57% vs. 20%, *P* = 0.16). Regarding treatment history, the anchor drug contained in ART1 was more frequently a non-nucleoside reverse transcriptase inhibitor (NNRTI) in SN than in SP (50% vs. 6%, *P* = 0.03), while current ART regimens did not differ between groups.

SN had significantly higher nadir CD4 T-cell counts since ART1 initiation than SP (1050 vs 604 cells/μL, *P* = 0.03). Consistent with this finding, current CD4 T-cell counts and percentages were higher in SN than in SP, although these differences were not significant (Figure 2A-B). The CD4 T-lymphocyte compartment tended to be slightly less differentiated in SN than in SP, as shown by lower effector and PD-1^+^ CD4 T-lymphocyte percentages (Figure 2C). The differentiation status of the CD8 T-lymphocyte compartment did not differ between SN and SP (Figure 2D). These findings support that, among early-treated children with sustained HIV suppression, negative HIV serology associated with higher nadir CD4 T-cell counts and a modest trend toward a less differentiated and less exhausted CD4 T-lymphocyte compartment at the time of the study.

**Figure 2.**
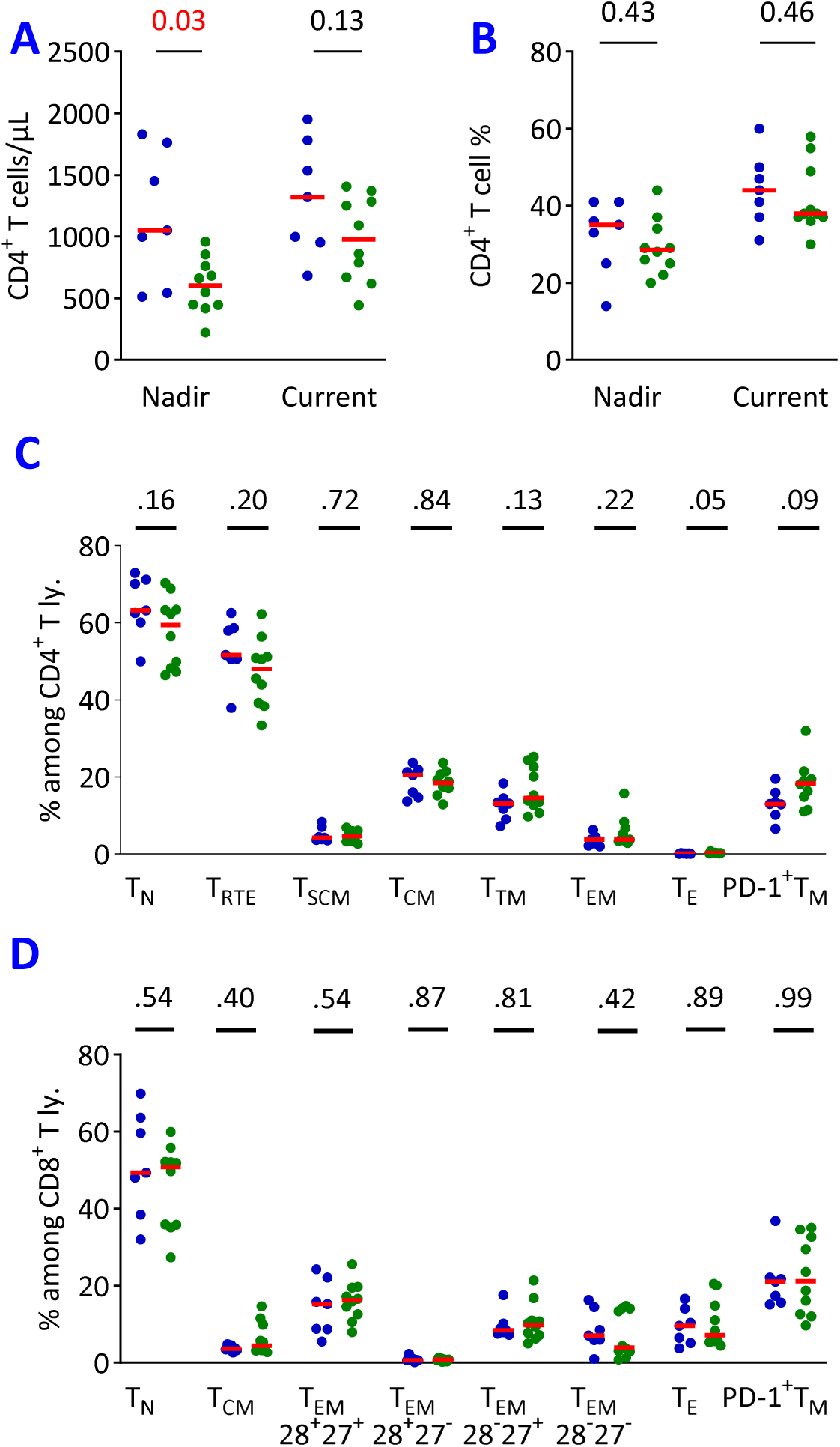
Higher nadir CD4 T cell-count associated with negative HIV serology. In the 17 early-treated children with sustained HIV suppression, we compared CD4 and CD8 T lymphocytes between SN and SP. A. CD4 T-cell counts at nadir after ART1 initiation and at the study visit. B. CD4 T-cell percentages at nadir after ART1 initiation and CD4 T-cell percentage at the study visit. C. We analyzed CD4 T lymphocyte subsets in fresh blood, including naive (T_N_, CD45RA^+^CCR7^+^), recent thymic emigrant (T_RTE_, CD45RA^+^CCR7^+^CD31^+^), stem cell memory (T_SCM_, CD45RA^+^CCR7^+^CD95^+^), central memory (T_CM_, CD45RA^-^CCR7^+^), transitional memory (T_TM_, CD45RA^-^CCR7^-^CD27^+^), effector memory (T_EM_, CD45RA^-^CCR7^-^CD27^-^), effector (T_E_, CD45RA^+^CCR7^-^), and PD-1^+^ memory (T_M_, CD45RA^-^ CCR7^+/-^) subsets. D. We analyzed CD8 T lymphocyte subsets in fresh blood, including naive (T_N_, CD45RA^+^CCR7^+^), central memory (T_CM_, CD45RA^-^CCR7^+^), effector memory (T_EM_, CD45RA^-^CCR7^-^) expressing or not CD27 and CD28, effector (T_E_, CD45RA^+^CCR7^-^), and PD-1^+^ memory (T_M_, CD45RA^-^CCR7^+/-^) subsets. Red bars indicate medians and *P* values above graphs correspond to Mann-Whitney tests.

### HIV-specific B and T lymphocytes were infrequently detected regardless of HIV serology

Next, we assessed whether SN and SP participants had detectable HIV-specific adaptive immune responses beyond plasma antibodies. Virus-specific B lymphocytes are heterogeneous and comprise two major subsets. Short-lived immunoglobulin (Ig)-producing plasmablasts produce the antibodies detected in plasma samples. Long-lived resting memory B lymphocytes do not produce Ig but can differentiate into Ig-producing cells upon restimulation [22]. We therefore investigated the presence of circulating HIV-specific memory B lymphocytes in the 4 SN and 5 SP participants for whom samples were available. We used the TLR7/8 ligand combined with the cytokine IL-2 to stimulate the differentiation of peripheral blood memory B lymphocytes into plasma cells [23,24]. After 7 days of culture, CD19⁺ B lymphocytes represented 50 to 80% of viable lymphocytes (Figure 3A). Among them, 50 to 70% were CD19⁺CD21ˡᵒCD27ʰⁱCD38ʰⁱ plasmablasts (Figure 3B). Culture supernatants contained IgG at concentrations ranging from 20 to 40 µg/mL, except for one SP participant whose B lymphocytes expanded but produced 10-fold less IgG than the other participants (Figure 3C). We detected IgG binding to recombinant p24, gp140, and gp41 proteins in cultures from one SN and IgG binding to p24 in cultures from one SP (Figure 3D), demonstrating the presence of circulating HIV-specific memory B lymphocytes. Addition of IL-6, a B-cell-stimulating cytokine, to the cultures did not impact plasmablast expansion or IgG production (S2-1 Figure), suggesting no defect in the lymphocytes and monocytes that produce this cytokine. Thus, *in vitro* expansion of memory B lymphocytes did not reveal any defect associated with negative HIV serology. Although rarely detected, HIV-specific B lymphocytes were present in both SN and SP.

**Figure 3.**
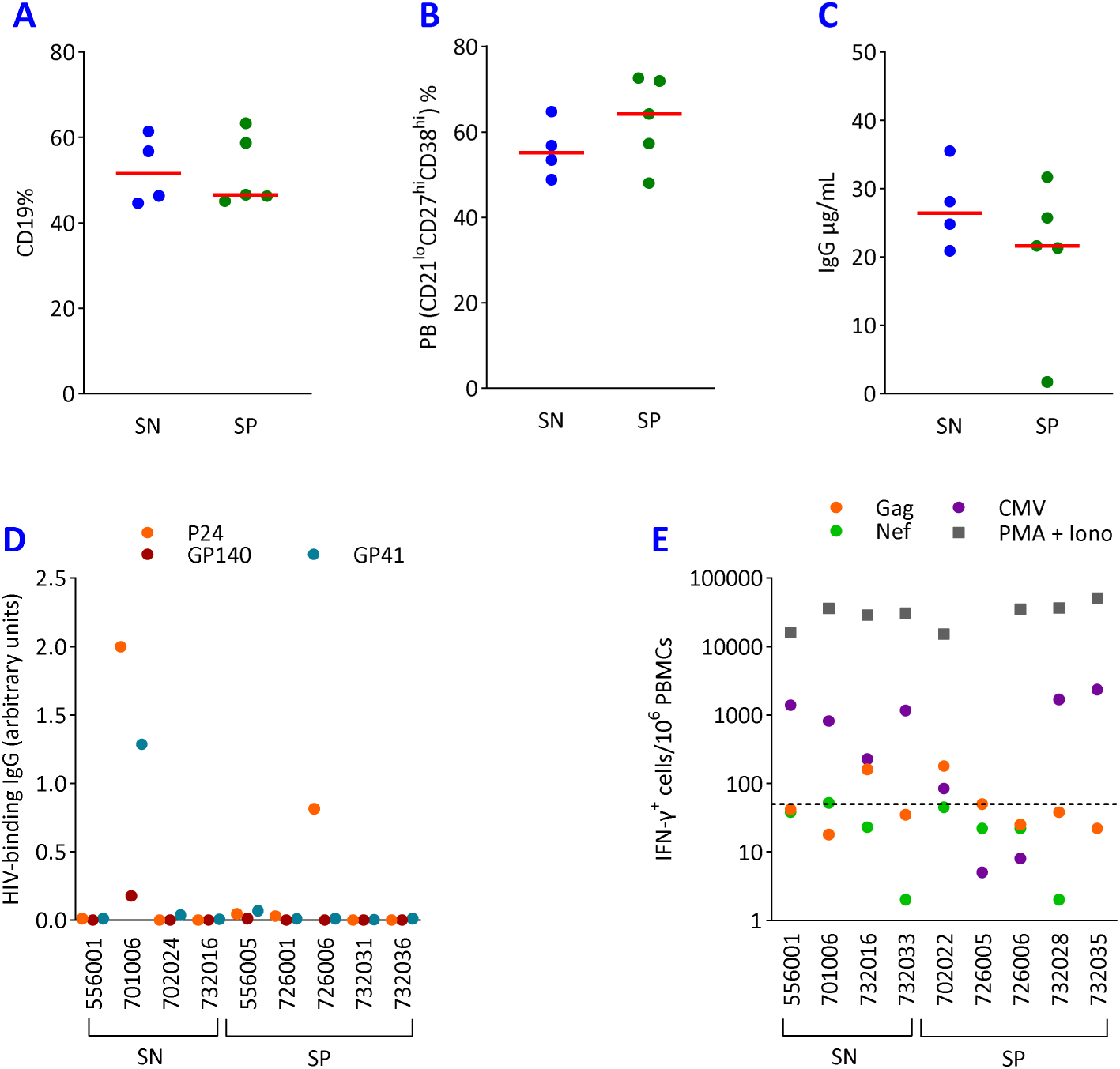
Both SN and SP participants harbored HIV-specific B and T lymphocytes. In the early-treated children with sustained HIV suppression, we compared HIV-specific B and T lymphocytes between SN and SP. Due to limited sample availability, we performed B- and T-cell assays on distinct participants (panels D and E). A to D. We thawed PBMCs, stimulated them with the TLR7/8 agonist R848, and cultured them with IL-2. We quantified by flow cytometry the percentages of B lymphocytes among viable cells (CD19+, panel A) and plasmablasts (CD19^+^CD21^lo^CD27^hi^CD38^hi^, panel B) among B lymphocytes at day 7. At day 12, we collected culture supernatants to quantify total (panel C) and HIV-specific IgG (panel D) by ELISA. E. We stimulated freshly isolated PBMCs with peptide pools or a mitogen and quantified IFN-γ production by ELISpot. The figure shows the frequency of IFN-γ-producing cells / 10^6^ PBMCs in response to peptide pools (HIV-1 Gag, HIV-1 Nef and HCMV pp65) and PMA / Ionomycin in nine children (4 SN, 5 SP). The dotted line indicates the positivity threshold (50 IFN-γ-producing cells/10^6^ PBMCs).

We then investigated HIV-specific T lymphocytes. We tested fresh PBMCs from nine children (4 SN and 5 SP) using an ELISPOT assay. We detected IFN-γ production in response to HIV-1 Gag and/or Nef peptide pools in 2 SN and 2 SP, demonstrating the presence of HIV-specific T lymphocytes in children without detectable HIV-specific antibodies (Figure 3E). To phenotype responding lymphocytes and expand the range of detected cytokines, we used an intracellular cytokine cytometry (ICC) assay on frozen PBMCs. We did not detect HIV-specific T lymphocytes participants analyzed using the ICC assay (S2-2 Figure), whose detection threshold was slightly higher than that of the ELISPOT assay, as described [25]. Both ELISPOT and ICC assays showed that T-cell responses to positive controls (HCMV peptides and mitogens) reached similar magnitudes in both groups (Figure 3E and S2-2 Figure). Thus, we detected HIV-specific T lymphocytes at low frequency regardless of HIV serology.

### Negative HIV serology associated with higher frequencies of circulating CXCR5^+^ CD8 T lymphocytes but not with the B-CD4 T lymphocyte axis

We next focused on the B–CXCR5^+^ CD4 T lymphocyte axis, as these lymphocyte subsets interact in lymph node germinal centers (GCs) to promote antibody production. We observed no differences between SN and SP in total B lymphocyte levels or their major subsets (Figure 4A). SN participants had lower plasma IgM levels than SP participants (0.39 vs 0.73 mg/mL, *P* = 0.01), but we detected other immunoglobulin isotypes at similar levels in both groups (Figure 4B). We quantified plasma molecules involved in B and T_FH_ differentiation, function, and migration (CXCL13, BAFF, APRIL, TGF-β, IL-6, and IL-21) and found comparable levels in plasma samples from SN and SP (Figure 4C). We detected CXCR5⁺CD4 T lymphocyte subsets defined by CXCR3 coexpression on T_CM_ and T_EM_, or by the PD-1⁺CXCR5⁺CXCR3⁻ phenotype, at similar levels in SN and SP (Figure 4D-E). Thus, our analysis of the B–CXCR5⁺T lymphocyte axis revealed no association with negative HIV serology.

**Figure 4.**
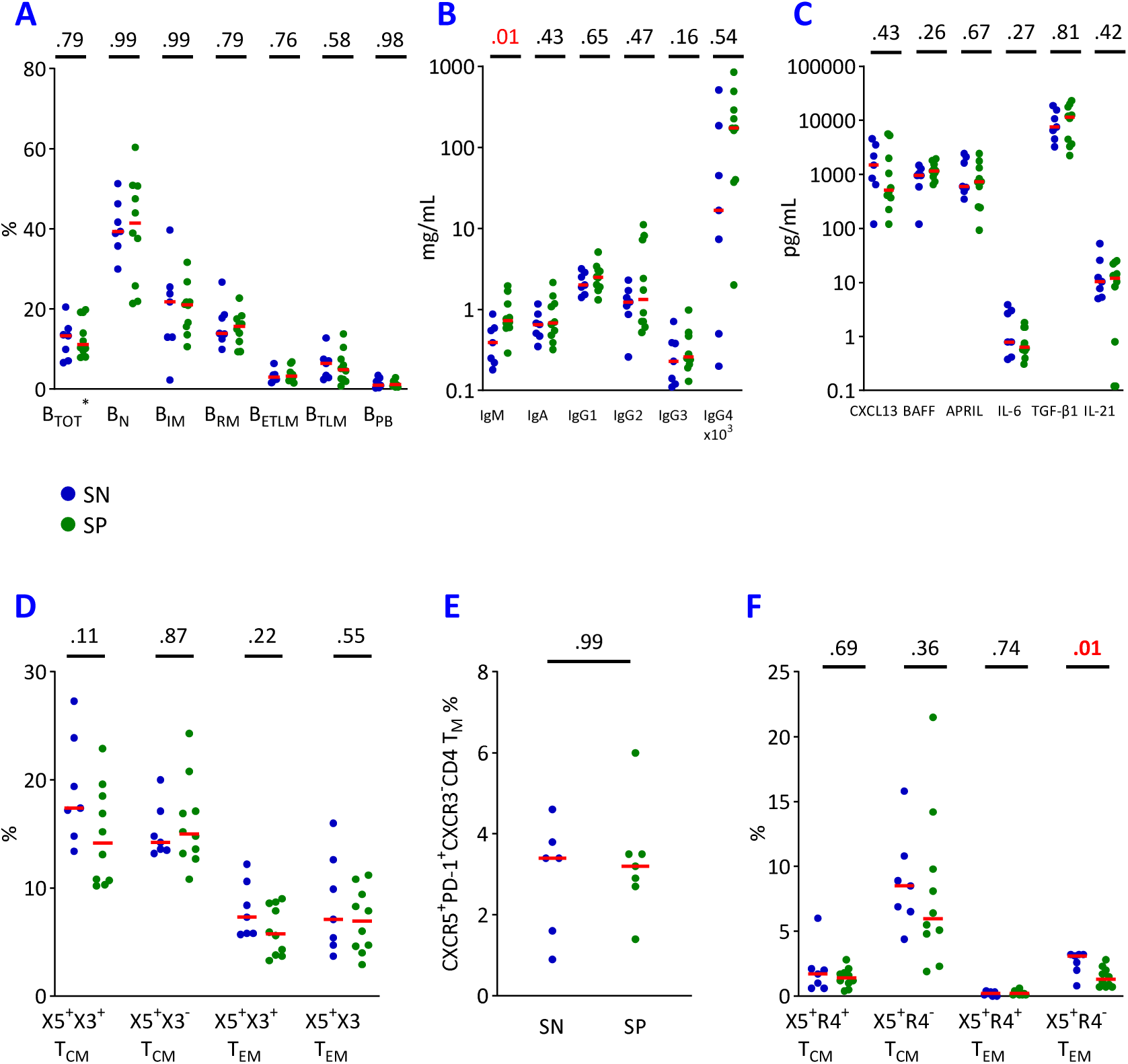
Negative HIV serology was associated with CXCR5+CD8 TEM lymphocytes but not with B and CD4 TFH lymphocytes. In the 17 early-treated children with sustained HIV suppression, we compared lymphocytes and soluble molecules involved in humoral immunity between SN and SP. A. Frequencies of total B lymphocytes (B_TOT_, CD19^+^, expressed as % among lymphocytes) and their subsets in fresh blood: naive (B_N_, CD10^-^CD21^+^CD27^-^), immature (B_IM_, CD10^+^CD21^+^CD27^-^), resting memory (B_RM_, CD10^-^CD21^+^CD27^+^), exhausted tissue-like memory (B_ETLM_, CD10^-^CD21^-^CD27^+^), tissue-like memory (B_TLM_, CD20^hi^CD10^-^CD21^lo^CD27^+^), and plasmablasts (B_PB_, CD20^-^CD27^+^CD38^hi^); % of CD19+ B lymphocytes). B. Plasma immunoglobulin concentrations, mg/mL. C. Plasma analytes related to B and T_FH_ differentiation, function, and migration, pg/mL. D. Frequencies of CXCR5^+^ cells expressing or not expressing CXCR3 among central memory (T_CM_, CD45RA^-^CCR7^+^) and effector memory (T_EM_, CD45RA^-^CCR7^-^) CD4 T lymphocytes, % among T_CM_ and T_EM_, respectively. E. Frequencies of T_FH_ defined as CXCR5^+^PD-1^+^CXCR3^-^ memory (CD45RA^-^CCR7^+/-^) CD4 T lymphocytes, % of memory CD4 T lymphocytes. F. Frequencies of CXCR5^+^ CD8 T_CM_ and T_EM_ lymphocytes coexpressing or not CCR4 among, % of T_CM_ and T_EM_, respectively. Red bars indicate medians and *P* values above graphs correspond to Mann-Whitney tests.

Besides being the site of antibody generation, GCs are a major site of HIV replication, which some CD8 T lymphocyte subsets reduce. We analyzed circulating CXCR5⁺ CD8 T lymphocytes based on CCR4 chemokine receptor coexpression on T_CM_ and T_EM_. SN participants had significantly higher CXCR5⁺CCR4⁻CD8 T_EM_ levels than SP participants (3.1 vs 1.3%, *P* = 0.01, Figure 4F). Studies have associated NK lymphocytes with HIV control in lymph nodes, but we found similar levels of blood NK cell subsets and NK receptor expression in both SN and SP (S2-3 Figure). Thus, among B and CXCR5^+^ T lymphocytes, only CCR4^-^ CD8 T_EM_ associated with negative HIV serology.

### Negative HIV serology associated with higher expression of costimulatory molecules on myeloid cells

Finally, we tested whether negative HIV serology associated with markers of T lymphocyte and myeloid cell-related activation, and plasma markers of immune activation, inflammation, and gut permeability. We found no differences between SN and SP in levels of activated HLA-DR⁺CD38⁺ memory T lymphocytes (CD4 and CD8 T_M_), regulatory CD4 T lymphocytes (T_REG_), or related cytokines/chemokines (Figures 5A and 5B). Similarly, the percentages of monocyte and dendritic cell (DC) subsets among mononuclear cells did not differ between groups (Figure 5C). However, CD14⁺CD16⁻ classical, CD14ˡᵒCD16⁺ nonclassical monocytes, and myeloid dendritic cells (mDCs) from SN expressed higher levels of the CD86 costimulatory molecule than those from SP (Figure 5D). Plasmacytoid dendritic cells (pDCs) from SN expressed higher levels of the coinhibitory PD-L1 molecule than those from SP (Figure 5E). In contrast, HLA-DR expression on myeloid cells (Figure 5F), plasma markers related to myeloid cells (Figure 5G), and those related to inflammation and gut permeability (Figure 5H) showed no differences between SN and SP participants. These findings identify higher expression of CD86 and PD-L1 on myeloid cells as a distinctive feature of negative HIV serology.

**Figure 5.**
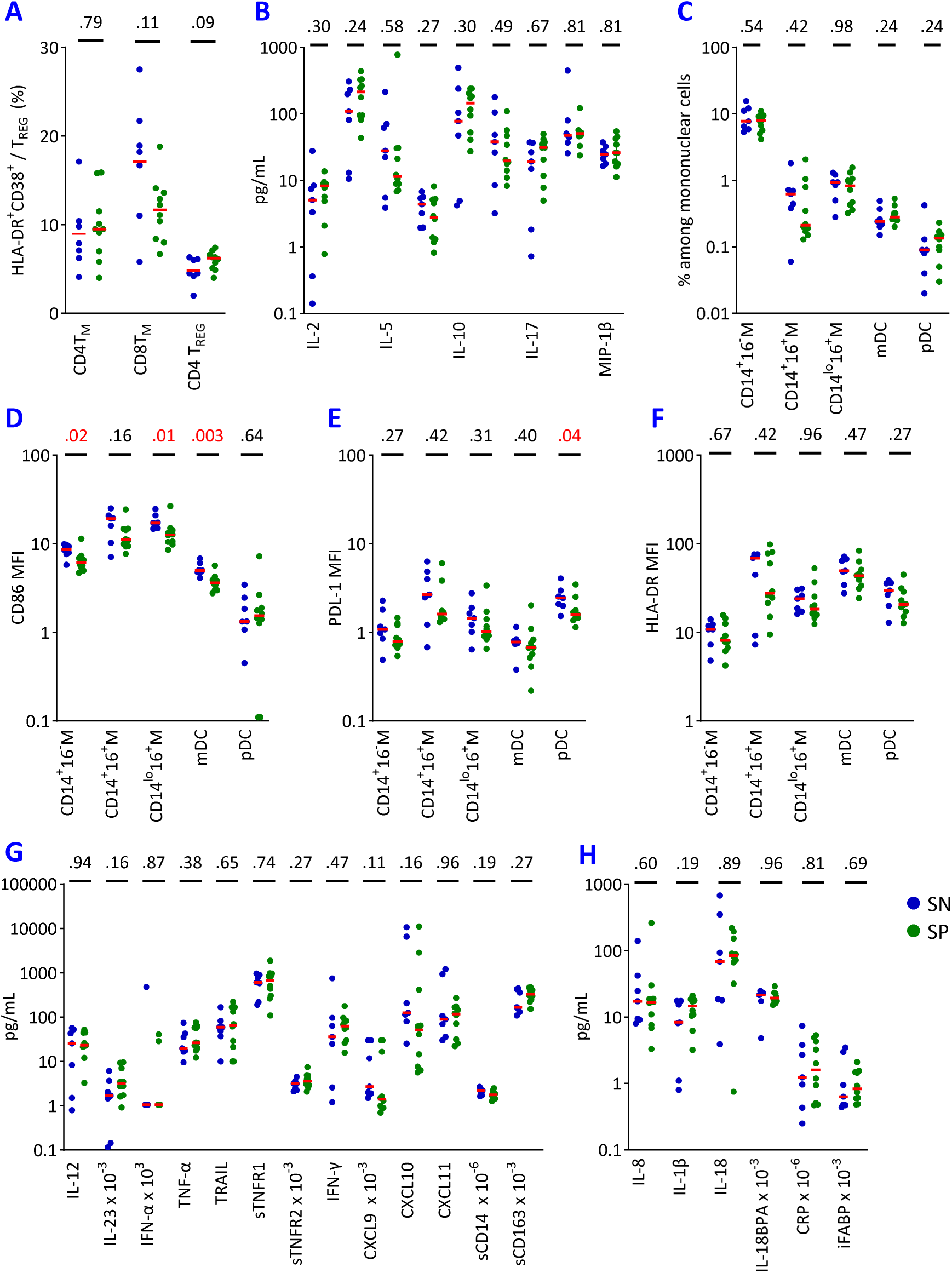
High costimulatory/coinhibitory molecule expression on monocytes and dendritic cells associated with negative HIV serology, unlike immune activation or inflammation markers. In the 17 early-treated children with sustained HIV suppression, we compared cells and soluble molecules related to immune activation, myeloid compartment and inflammation between SN and SP A. Frequencies of activated CD4 and CD8 memory T lymphocytes (HLA-DR^+^CD38^+^ non-CD45RA^+^CCR7^+^, % of CD4 and CD8 T_M_ lymphocytes) and regulatory CD4 T lymphocytes (T_REG_, CD25^hi^CD127^lo^, % of CD4 T lymphocytes). B. Plasma concentrations of T-cell related cytokines and chemokines (pg/mL). C. Frequencies of myeloid cell subsets: classical CD14^+^CD16^-^, intermediate CD14^+^CD16^+^ and activated CD14^-/lo^CD16^+^ monocytes, mDCs and pDCs (% of mononuclear cells). D to F. We quantified expression levels of CD86/PD-L1/HLA-DR on the five subsets of myeloid cells (mean fluorescence intensity, MFI). G. Plasma concentrations of molecules related to innate immunity and myeloid cells (pg/mL). H. Plasma concentrations of inflammation-related molecules and the gut permeability marker, intestinal fatty acid-binding protein (pg/mL). Red bars indicate medians and *P* values above graphs correspond to Mann-Whitney tests.

### Negative HIV serology associated with past and current immune parameters

We analyzed the correlations among variables which associated or tended to associate with negative HIV serology in previous analyses, i.e., nadir CD4 T-cell count, age at viral suppression, plasma IgM levels, CXCR5⁺CCR4⁻ CD8 T_EM_ percentages, expression levels of CD86 on CD14^+^CD16^-^ monocytes, CD14^lo^CD16^+^ monocytes, mDCs, and the expression level of PD-L1 on pDCs (Figure 6A and S2-4 Figure). Nadir CD4 T-cell values did not correlate with any other variables. Plasma IgM correlated positively with age at viral suppression and negatively with CXCR5⁺CCR4⁻ CD8 T_EM_ percentages. CD86 and PD-L1 expression levels on myeloid cells correlated with each other, and CD86 expression correlated with CXCR5⁺CCR4⁻ CD8 T_EM_ percentages.

**Figure 6.**
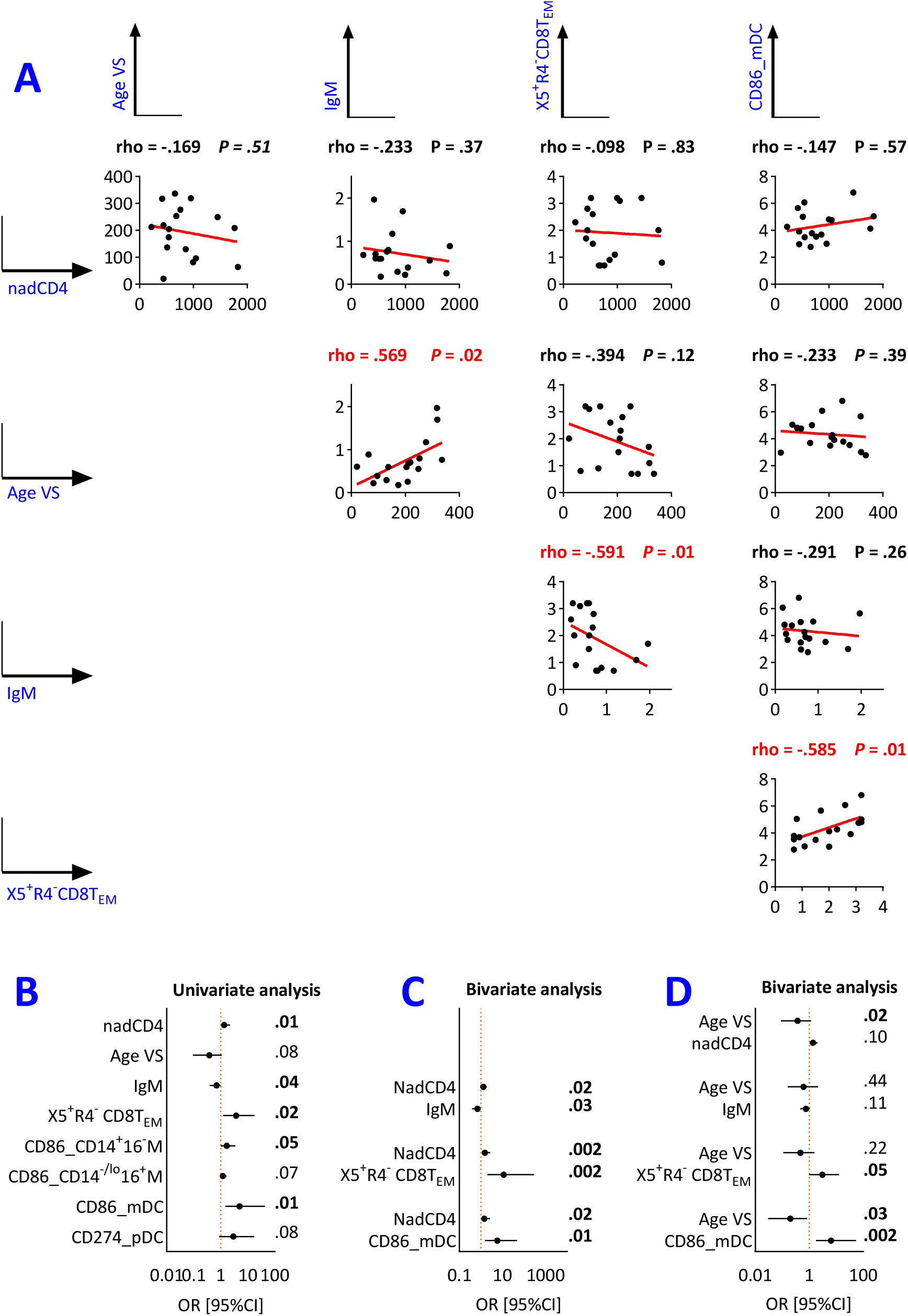
Correlations and logistic regression analysis of variables associated with negative HIV serology. A. We used the Spearman test to assess correlations between continuous variables associated with negative HIV serology (Age VS, Age at first HIV RNA < 400 copies/mL expressed, days; IgM, µg/mL; X5^+^R4^-^CD8T_EM_, CXCR5^+^CCR4^-^ CD8T_EM_, %; CD86_mDC, CD86 MFI on mDCs ; nadCD4, Nadir CD4, cells/µL). Coefficients and *P* values are indicated on the graphs. Red characters denote *P* values < 0.05. B to D. We used Firth’s penalized logistic regression with added covariates to analyze negative HIV serology, including non-collinear variables. Odds ratios with 95% confidence intervals and *P* values are shown. Bold characters denote *P* values < 0.05. B. univariate analysis. C and D. Bivariate analysis including nadir CD4 T-cell counts or age at HIV RNA < 400 copies/mL, respectively. We calculated odds ratios per 100 cells/µL for CD4 counts, per 100 days for age, per 10 mg/mL for IgM, per 1% for CD8, and per 1 unit of MFI for CD86 and PD-L1 expression.

We then used Firth’s logistic regression to identify noncollinear variables associated with negative HIV serology [19]. In univariate analysis, negative HIV serology significantly associated with higher nadir CD4 T-cell count, lower plasma IgM levels, higher CXCR5⁺CCR4⁻ CD8 T_EM_ percentages, and higher CD86 expression on mDCs (Figure 6B). We then built two-variable models, each pairing one variable related to past HIV history (nadir CD4 count or age at viral suppression) with one immunological variable (Figure 6B). In these models, negative HIV serology independently associated with nadir CD4 T-cell counts and with each of the three immunological variables. After adjustment for age at viral suppression, negative HIV serology remained significantly associated with CXCR5⁺CCR4⁻ CD8 T_EM_ percentages, and CD86 expression on mDCs, but not with IgM levels. Notably, negative HIV serology remained significantly associated with nadir CD4 T-cell counts after this adjustment. Models including more than two independent variables could not be reliably fitted, most likely due to the limited sample size. Taken together, and with caution given the number of participants, these analyses support that negative HIV serology may independently associate with both past HIV-related immunosuppression and current phenotypic markers (CXCR5⁺CCR4⁻ CD8 T_EM_ percentages or CD86 expression on mDCs).

## Discussion

We report a strong association between negative HIV serology and both early ART initiation and reduced lifetime exposure to HIV replication among children and adolescents infected with HIV in the perinatal period who initiated ART early (< 6 months of age) or late (>24 months of age) during childhood. Nevertheless, fewer than half of children who achieved viral suppression at a young age and had low HIV DNA levels showed negative HIV serology. For early-treated children with sustained viral suppression, we identified several parameters significantly associated with negative HIV serology: higher nadir CD4 T-cell counts after ART initiation, lower plasma IgM, higher CXCR5⁺CCR4⁻ CD8 T_EM_ percentages, higher CD86 expression on monocytes and mDCs, and higher expression of PD-L1 on pDCs. Additionally, younger age at viral suppression approached significance for an association with negative HIV serology.

Other studies examining children with varying levels of HIV replication and ART exposure consistently show robust associations between negative HIV serology, early ART initiation and low viral reservoir size [6,26–29]. However, when cohorts comprise only children with early and sustained HIV suppression, participants have similarly low viral exposure and these associations are inconsistent: HIV-specific antibody levels vary considerably [6,30] and show no correlation with HIV DNA levels [13,31–33]. We recently described negative HIV serology in an adolescent with perinatal HIV-1 infection who initiated ART at one month of age but had sustained viral replication until four years of age [34]. Collectively, these data indicate that limited HIV antigen exposure is necessary but insufficient to explain negative HIV serology.

To identify factors associated with negative HIV serology, we selected children with low HIV antigen exposure and compared those with and without detectable HIV serological responses. SN and SP showed no significant differences in most virus-related variables. Because HIV replication potently drives naive CD8 T-lymphocyte recruitment into the memory compartment, we examined T-lymphocyte phenotypes. Both groups exhibited similar proportions of naive and memory CD8 T lymphocytes, confirming comparable HIV-induced immune changes. Thus, our selection criteria adequately controlled for HIV exposure when investigating immune factors related to negative HIV serology.

After initiating ART1, SN tended to achieve viral suppression earlier than SP. Although this association did not reach significance, it warrants discussion. Being younger during the period of exposure to HIV antigens may reduce anti-HIV antibody responses through several mechanisms: impaired B-lymphocyte function [35], maternal HIV-specific antibodies that interfere with infant B-lymphocyte stimulation by HIV antigens, and reduced cumulative exposure to these antigens [36,37]. We found that negative HIV serology did not associate with past HIV replication levels or current reservoir size, suggesting that HIV antigen load was not the underlying mechanism. Rather, immune development and/or maternal HIV-specific antibody levels may explain the relationship between age at viral suppression and HIV serology in early-treated children.

We found no association between negative HIV serology and the viro-immunological profile we previously described in the early-treated children, i.e., low HIV DNA levels and less differentiated CD8 T lymphocytes [14,15]. Instead, SN exhibited higher nadir CD4 T-cell counts, lower IgM levels, and a trend toward younger age at viral suppression compared to SP. These findings suggest that SN would have experienced less aggressive disease progression than SP in the absence of ART. Negative HIV serology was more frequent in children who received NNRTI-based regimen as first ART than in those who received a protease inhibitor (PI)-based regimen. NNRTIs have been suggested to be more potent than PIs in reducing residual viral transcription and viral reservoirs [38,39]. In addition, the two classes of antiretroviral molecules have distinct effects on antigenic stimulation because NNRTIs prevent HIV protein expression whereas PIs do not [40].

Follicular CD8 T (fCD8) lymphocytes express CXCR5 but not CCR7, enabling their homing to lymphoid follicles—key sites of antibody production and sanctuaries for HIV replication. These lymphocytes exert antiviral activity against HIV/SIV replicating within follicles, predominantly in untreated controllers [41–43]. In HIV-infected adults, peripheral blood fCD8 frequencies correlate inversely with plasma HIV RNA [44]. SN had significantly higher levels of CCR4^-^CXCR5^+^ among CD8 T_EM_ lymphocytes than SP. This difference was specific to the T_EM_ lymphocytes, which do not express CCR7, and was not observed for CXCR5^+^CD8 T_CM_ lymphocytes, which express CCR7 and do not home to follicles. In children, HIV infection reduces fCD8 frequency in association with CD4 T-lymphocyte depletion and elevated HIV RNA levels [45], and circulating fCD8 T-lymphocyte frequency correlates positively with neutralization breadth [46]. The elevated circulating fCD8 T-lymphocyte frequencies in SN suggest that these lymphocytes may reduce HIV replication within lymphoid follicles, thereby limiting B-lymphocyte stimulation by viral antigens. Furthermore, their antiviral capacity may confer greater potential for immune control should HIV replication resume.

We identified phenotypic properties of myeloid cells associated with negative HIV serology. Costimulatory molecules such as CD86 ensure optimal antigen presentation to T lymphocytes and their effective activation. HIV infection decreases CD86 expression, while ART preserves it [47]. PD-L1, a coinhibitory molecule, dampens antiviral lymphocyte activity but may prevent the deleterious effects of excessive immune activation [48]. The crosstalk between these regulatory molecules fine-tunes lymphocyte responses [49]. Consistent with this immunoregulatory framework, we observed differences in both the levels of CD86 expression on myeloid cells and the frequencies of fCD8 T-lymphocytes between SN and SP, and found correlations between CD86 expression on myeloid cells and fCD8 T lymphocyte frequency. These findings suggest two mechanisms that are not mutually exclusive: efficient cell-mediated immunity in lymph nodes may reduce HIV replication and subsequent B-lymphocyte antigenic stimulation, or preserved fCD8 lymphocytes and myeloid cells may reflect limited HIV-induced alterations in lymphoid tissues.

Negative HIV serology has been proposed as a clinical marker to identify patients with low HIV reservoirs eligible for remission strategies. We therefore verified that undetectable HIV antibodies do not reflect immune defects. Our phenotypic study and plasma markers found no evidence that B or CD4 T_FH_ lymphocyte deficits underlie negative HIV serology. Furthermore, the absence of HIV antibodies by diagnostic EIA does not reflect a globally tolerant state characterized by the absence of HIV-specific T and B lymphocytes. Using three functional assays, we detected low-frequency, low-magnitude HIV-specific adaptive immune responses that were present in both SN and SP. These data demonstrate that the brief period of HIV replication in infancy can generate sufficient antigenic stimulation to induce HIV-specific T and B lymphocytes, at least in some children. Thus, HIV-specific antibodies may be undetectable despite the presence of HIV-specific B and T lymphocytes, as other studies have reported [4,30,50].

Beyond evaluation of the putative immune defects, our systematic analysis of B-cell-related markers revealed a further distinction between groups. Among B-cell-related molecules, only IgM levels differed between SN and SP, with lower levels in SN. Polyclonal hypergammaglobulinemia represents one of the earliest markers of HIV-induced immune activation [51–53]. At birth, HIV-infected children show expansion of transitional B lymphocytes and plasmablasts, and ART normalizes this pattern within weeks [13]. Lower IgM levels in SN may reflect reduced B-cell activation before ART-induced viral suppression and may explain their weaker antibody response to HIV antigens. However, another study of early-treated children with viral suppression found similar IgM levels between children with and without anti-HIV antibodies [30]. Under ART, some proviruses remain transcriptionally active and viral proteins may stimulate antibody production and induce immune activation [54,55]. This mechanism is unlikely to explain negative HIV serology in our study, as negative HIV serology did not associate with plasma p24 antigen or classical markers of HIV-induced immune activation [53,56,57]. A study of adults treated during acute infection found no association between negative HIV serology and immune activation on ART, although such an association was present before ART initiation [58]. Thus, among children with low HIV replication exposure, current immune activation markers did not associate with negative HIV serology. Whether current IgM levels reflect pre-ART B-lymphocyte activation and subsequent HIV-specific antibody induction warrants further investigation.

Our study has several limitations. First, we observed negative HIV serology in 5-to-12-year-old children, although physiological events during the first months of life determined, at least in part, this outcome. Although studies at the time of early life HIV antigen exposure would better decipher the mechanisms underlying antibody responses, cross-sectional studies performed years later remain important for identifying biomarkers useful for HIV remission strategies. Second, we miss some data pertaining to HIV history. HIV RNA and CD4 T-cell counts were systematically assessed once ART began, but these laboratory measures were missing for some children in the pre-ART period, which prevented precise estimation of early life HIV replication exposure. We also lacked CD4 T-cell measures at ART initiation for some children—information that would help interpret the nadir CD4 T-cell counts they reached under ART. For some participants, we could not determine whether HIV transmission occurred in utero or peripartum. Consequently, we did not assess the impact of exposure to HIV during fetal life on the generation of antibodies against HIV. Third, our study is exploratory and data were obtained from a small number of children. Fourth, exposure to infectious agents other than HIV and genetic background vary across geographical contexts [59]. Thus, studies conducted in different settings may report different associations. Fifth, we derived most immune parameters from a pre-specified phenotypic study designed to cover major cell subsets, while we performed functional assays on frozen cells. Limited blood sample volume prevented functional characterization of all cell subsets of interest. We chose to assess B-lymphocyte function *in vitro* because these cells produce HIV-specific antibodies. We acknowledge that functional studies of CD8 T lymphocytes and CD86 expression on myeloid cells would have strengthened our phenotypic findings.

In conclusion, despite the major role of viral replication in inducing HIV antibodies, the underlying complexity of immune responses must be considered. We identified associations between negative HIV serology and higher nadir CD4 counts, lower IgM levels, and higher fCD8 T-lymphocyte frequencies—the latter have potent antiviral activity and are capable of migrating to lymph node follicles. Additionally, we found no evidence of defects or dysregulation in the B-T_FH_ lymphocyte axis. This exploratory study supports negative HIV serology as a valuable clinical marker in HIV remission strategies, as it associates with fCD8 T-lymphocyte frequency and less aggressive disease progression.

## Acknowledgements

We thank the children and adolescents who participated in the study and their families. We thank colleagues from the cytometry platform, the UTechS Single cells biomarkers, and Viral Reservoirs and Immune Control unit at Institut Pasteur for their technical assistance. The peptide pools were obtained through the AIDS Reagent Program, Division of AIDS, NIAID, NIH.

## The ANRS CLEAC Study group

Hôpital Armand Trousseau, Paris: Mary-France Courcoux, Catherine Dollfus, Marie-Dominique Tabone, Geneviève Vaudre; Hôpital Bicêtre, Le Kremlin-Bicêtre: Corinne Fourcade, Josiane Warsazawski, Jérôme Lechenadec, Olivia Dialla, Laura Nailler, Lamya Ait-Si-Selmi, Isabelle Leymarie, Thierry Wack, Alexandre Hoctin, Razika Feraon-Nanache; Centre hospitalier intercommunal de Créteil: Isabelle Hau; Hôpital Delafontaine, Saint-Denis: Cécile Gakobwa; Hôpital Necker-Enfants Malades, Paris: Véronique Avettand-Fenoël, Stéphane Blanche, Marine Fillion, Pierre Frange, Nizar Mahlaoui, Adeline Mélard, Elise Gardenniet, Florence Veber, Marie-Christine Mourey; Maternité Port-Royal, Paris: Valérie Marcou ; Hôpital Robert Debré, Paris: Albert Faye, Martine Lévine, Sandrine Richard; Pitié-Salpêtrière, Paris: Brigitte Autran, Assia Samri, Mariama Diallo; Hôpital Saint-Louis: Sophie Caillat-Zucman, Kahina Amokrane, Rayna Ivanova-Derin, Centre hospitalier intercommunal de Villeneuve-Saint-Georges: Anne Chacé; Institut Pasteur Paris: Florence Buseyne, Thomas Montange, Damien Batalie, Ingrid Fert, Asier Saez-Cirion, Valérie Monceaux, Daniel Scott-Algara; ANRS-MIE: Lucie Marchand, Delphine Lebrasseur, Axel Levier. ANRS (France Recherche Nord et Sud SIDA-HIV Hépatites) is the sponsor of the study.

