## Supplemental materials, methods and data for "Negative HIV serology in perinatally infected children after early treatment and viral suppression: an exploratory analysis of immune correlates"

### Frange et al. Supplemental information

### Supplemental information – Part 1: immunological assays performed for the ANRS-EP59-CLEAC study

#### Quantification of plasma analytes

We collected plasma from EDTA-anticoagulated blood and stored samples at -80°C. We quantified analytes according to the manufacturer's instructions. For Luminex and Milliplex assays, we prepared samples using DropArray™ 96 plates (Curiox) and acquired data on a BioPlex 200 (Biorad). For SIMOA assays, we acquired data on an HD-X analyzer (Quanterix). The table presents the reference, quantification threshold, percentages of samples below the quantification threshold, and percentages of samples with extrapolated values.

**Table S1-1. Assays for plasma analyte quantification**

| Analyte | Assay type[1] | Manufac-turer[1] | Cat N° | Quantification threshold | % (n) below detection threshold among 76 patients tested for the CLEAC study [% (n)] extrapolated |
|---|---|---|---|---|---|
| Soluble CD163 (sCD163) | ELISA | RDS | DY1607 | 156 pg/mL | 0.0 (0) |
| C-Réactive Protéine (CRP) | ELISA | RDS | DY1707 | 15.6 pg/mL | 0.0 (0) |
| soluble transferrin receptor (sTfR) | ELISA | RDS | DY2474 | 78.1 pg/mL | 0.0 (0) |
| Soluble CD14 (sCD14) | ELISA | RDS | DY383 | 62.5 pg/mL | 0.0 (0) |
| Transforming growth factor beta-1 (TGF-β1) | ELISA | RDS | DY240 | 31.3 pg/mL | 0.0 (0) |
| Lipopolysaccharide binding protein (LBP) | ELISA | RDS | DY870 | 0.78 ng/mL | 0.0 (0) |
| A proliferation-inducing ligand / TNFSF13 (APRIL) | ELISA | RDS | DY884B | 20 pg/mL | 0.0 (0) [6.7(5)] |
| IL-18 Binding Protein A (IL18BPA) | ELISA | RDS | DY119 | 4 pg/mL | 0.0 (0) [2.6 (2)] |
| Intestinal fatty acid binding protein (I-FABP) | ELISA | CS | HK406-02 | 180 pg/mL | 1.3 (1) |
| Endotoxin Core Antibodies (EndoCab-IgM) | ELISA | CS | HK504-IgM | 0.05 MMU/mL | [2.6 (2)] |
| Immunoglobulin M (IgM) | Multiplex | MM | HGAMMAG-301K | 3.43 ng/mL | 0.0 (0) |
| Immunoglobulin G1 (IgG1) | Multiplex | MM | HGAMMAG-301K | 13.72 ng/mL | 0.0 (0) |
| Immunoglobulin G2 (IgG2) | Multiplex | MM | HGAMMAG-301K | 123 µg/mL | 0.0 (0) [1.3(1)] |
| Immunoglobulin G3 (IgG3) | Multiplex | MM | HGAMMAG-301K | 33 µg/mL | 5.2 (4) [3.9(3)] |
| Immunoglobulin G4 (IgG4) | Multiplex | MM | HGAMMAG-301K | 0.2 µg/mL | 0.0 (0) [11.8(9)] |

| | | | | | |
|---|---|---|---|---|---|
| Immunoglobulin A (IgA) | Multiplex | MM | HGAMMAG-301K | 2.06 µg/mL | 0.0 (0) |
| Interleukin-18 (IL-18) | Multiplex | MM | HIL18MAG-66K | 1.5 pg/mL | 3.9 (3) [6.7(5)] |
| soluble gp130 (sgp130) | Multiplex | MM | HSCRMAG-32PMX14BK | 24 pg/mL | 0.0 (0) |
| soluble Interleukin-6 Receptor (sIL-6R) | Multiplex | MM | HSCRMAG-32PMX14BK | 12 pg/mL | 0.0 (0) |
| soluble Tumor Necrosis Factor Receptor I (sTNFRI) | Multiplex | RDS | LXSAHM-02 | 0.9 pg/mL | 7.9 (6) [9.2(7)] |
| soluble Tumor Necrosis Factor Receptor II (sTNFRII) | Multiplex | RDS | LXSAHM-02 | 12 pg/mL | [5.3(4)] |
| B cell activating factor (BAFF) | Multiplex | RDS | LXSAHM-01 | 0.24 ng/mL | 4.0 (3) |
| chemokine (C-X-C motif) ligand 9 (CXCL9) | ELISA | RDS | DY392 | 62.5 pg/mL | [10.5(8)] |
| chemokine (C-X-C motif) ligand 10 (CXCL10) | ELISA | RDS | DY06605 | 4 pg/mL | 4.0 (3) [1.3(1)] |
| chemokine (C-X-C motif) ligand 13 (CXCL13) | ELISA | RDS | DY801 | 12 pg/mL | 4.0 (3) [1.3(1)] |
| tumor-necrosis-factor related apoptosis inducing ligand (TRAIL) | Multiplex | RDS | LXSAHM-05 | 20 pg/mL | 23.7 (18) + [18.4 (14)] |
| Interferon-inducible T-cell alpha chemoattractant (ITAC/CXCL11) | Multiplex | MM | HSTCMAG-28PMX21BK | 2.4 pg/mL | 0.0 (0) |
| Fractalkine (CX3CL1) | Multiplex | MM | HSTCMAG-28PMX21BK | 77 pg/mL | 13.1 (10) |
| Monokine with inflammatory and chemokinetic properties (MIP-1a) | Multiplex | MM | HSTCMAG-28PMX21BK | 3.9 pg/mL | 0.0 (0) |
| Monokine with inflammatory and chemokinetic properties (MIP-1b) | Multiplex | MM | HSTCMAG-28PMX21BK | 2.9 pg/mL | 0.0 (0) |
| Monokine with inflammatory and chemokinetic properties (MIP-3a/CCL20) | Multiplex | MM | HSTCMAG-28PMX21BK | 2.4 pg/mL | 5.2 (4) |
| Granulocyte-macrophage colony-stimulating factor (GM-CSF) | Multiplex | MM | HSTCMAG-28PMX21BK | 1.22 pg/mL | 11.8 (9) |
| Interferon gamma (IFNg) | Multiplex | MM | HSTCMAG-28PMX21BK | 2 pg/mL | 7.9 (6) |
| Interleukin-2 (IL-2) | Multiplex | MM | HSTCMAG-28PMX21BK | 0.5 pg/mL | 0.0 (0) [23.7(18)] |
| Interleukin-17A (IL-17A) | Multiplex | MM | HSTCMAG-28PMX21BK | 0.73 pg/mL | 0.0 (0) [11.8(9)] |
| Interleukin-21 (IL-21) | Multiplex | MM | HSTCMAG-28PMX21BK | 0.24 pg/mL | 7.8 (6) [9.2(7)] |
| Tumor necrosis factor alpha (TNFa) | Multiplex | MM | HSTCMAG-28PMX21BK | 0.9 pg/mL | 0.0 (0) |
| Interleukin-13 (IL-13) | Multiplex | MM | HSTCMAG-28PMX21BK | 0.98 pg/mL | 7.9 (7) [6.7(5)] |
| Interleukin-4 (IL-4) | Multiplex | MM | HSTCMAG-28PMX21BK | 4 pg/mL | 9.2 (7) |
| Interleukin-5 (IL-5) | Multiplex | MM | HSTCMAG-28PMX21BK | 0.45 pg/mL | 5.2 (4) [1.3(1)] |
| Interleukin-1 beta (IL-1b) | Multiplex | MM | HSTCMAG-28PMX21BK | 0.49 pg/mL | 0.0 (0) [11.8(9)] |
| Interleukin-6 (IL-6) | ELISA | RDS | HS600B | 0.1 pg/mL | 0.0 (0) |
| Interleukin-7 (IL-7) | ELISA | RDS | HS750 | 0.3 pg/mL | 1.3 (1) |

| | | | | | |
|---|---|---|---|---|---|
| Interleukin-8 (IL-8) | Multiplex | MM | HSTCMAG-28PMX21BK | 0.3 pg/mL | 0.0 (0) |
| Interleukin-10 (IL-10) | Multiplex | MM | HSTCMAG-28PMX21BK | 0.5 pg/mL | 9.2 (7) |
| Interleukin-12 (IL-12p70) | Multiplex | MM | HSTCMAG-28PMX21BK | 0.5 pg/mL | [100(76)] |
| Interleukin-23 (IL-23) | Multiplex | MM | HSTCMAG-28PMX21BK | 32 pg/mL | 7.8 (6) |
| Interferon alpha (IFNa) | SIMOA | Qx | 100860 | 0.021 pg/mL | 79.5(31/39) |
| HIV p24 antigen | SIMOA | Qx | 102215 | 0.024 pg/mL | 65.8 (25/38) |

[1] ELISA, enzyme-linked immunoabsorbent assay; Multiplex, multiplex bead-based immunoassay; SIMOA, single molecule array.
[2] CS, Cliniscience; MM, Merck-Millipore ; Qx, Quanterix; RDS, R&D systems.

#### Phenotypic study of mononuclear cells performed on fresh blood samples

##### Staining conditions

We performed mononuclear cells phenotypic studies on fresh blood samples collected on EDTA tubes. We stained 100 µl or 200 µl of whole blood according to the frequency of analyzed cell subsets, with eight mixtures of antibodies in 5 mL tubes (Becton Dickinson, 352054) for 30 min at room temperature (RT) in the dark. We then added 3 mL of red blood cell lysis kit (eBioscience, 00-5333-57), mixed samples by low speed vortexing, and incubated the cells for 15 min at RT. We centrifuged the cells for 6 min at 500 x *g*, washed once in 2 mL PBS, and resuspended the pellet in 0.3 mL PBS and immediately analyzed stained samples. The staining conditions and cytometer's detectors are summarized in tables S1-2 and S1-3, respectively.

**Table S1-2: The eight sets of staining conditions for mononuclear cell subsets analysis**

| Tube name | Blood volume, µL | Antigen | Fluorochrome | Clone | Volume, µl | Manufacturer[1] | Cat N° |
|---|---|---|---|---|---|---|---|
| Tnai | 100 | CD45RA | BV421 | Hi100 | 1 | BD | 562885 |
| | | CD3 | Krome Orange | UCHT1 | 1 | BC | B00068 |
| | | CD57 | FITC | NK-1 | 5 | BD | 555619 |
| | | CD279 | PE | J105 | 1 | eB | 12-2799 |
| | | CD27 | ECD | 1A4CD27 | 1 | BC | B26603 |
| | | CCR7 | PC7 | 3D12 | 2 | BD | 560922 |
| | | CD28 | APC | CD28.2 | 1 | BC | B10244 |
| | | CD4 | Alexa 700 | RPA-T4 | 1 | BD | 561030 |
| | | CD8α | APC-Cy7 | RPA-T8 | 1 | BD | 557760 |
| Trte | 100 | CD45RA | BV421 | Hi100 | 1 | BD | 562885 |
| | | CD3 | Krome Orange | UCHT1 | 1 | BC | B00068 |
| | | CD31 | FITC | 5.6E | 5 | BC | IM1431U |
| | | CD95 | PE | DX2 | 5 | BD | 561976 |
| | | CD27 | ECD | 1A4CD27 | 1 | BC | B26603 |
| | | CCR7 | PC7 | 3D12 | 2 | BD | 560922 |
| | | CD28 | APC | CD28.2 | 1 | BC | B10244 |
| | | CD4 | Alexa 700 | RPA-T4 | 1 | BD | 561030 |
| | | CD62L | APC-H7 | DREG56 | 2 | BC | B26604 |
| Tact | 100 | CD45RA | BV421 | Hi100 | 1 | BD | 562885 |

| | | | | | | | |
|---|---|---|---|---|---|---|---|
| | | CD3 | Krome Orange | UCHT1 | 1 | BC | B00068 |
| | | HLA-DR | FITC | 9-49 | 2 | BC | 6603424 |
| | | CD127 | PE | R34.34 | 5 | BC | IM1980U |
| | | CD38 | ECD | LS198.4.3 | 1 | BC | A99022 |
| | | CCR7 | PC7 | 3D12 | 2 | BD | 560922 |
| | | CD25 | APC | 24212 | 1 | RDS | FAB1020A |
| | | CD4 | Alexa 700 | RPA-T4 | 1 | BD | 561030 |
| | | CD8α | APC-Cy7 | RPA-T8 | 1 | BD | 557760 |
| Tpol | 100 | CD45RA | BV421 | Hi100 | 1 | BD | 562885 |
| | | CD3 | Krome Orange | UCHT1 | 1 | BC | B00068 |
| | | CXCR5 | FITC | 51505 | 5 | RDS | FAB190F |
| | | CCR6 | PE | 11A9 | 1 | BD | 561019 |
| | | CCR4 | PE-CF594 | 1G1 | 5 | BD | 565391 |
| | | CD183 | PC5 | 1C6 | 2 | BD | 561731 |
| | | CCR7 | PC7 | 3D12 | 2 | BD | 560922 |
| | | CD4 | Alexa700 | RPA-T4 | 1 | BD | 561030 |
| | | CD8a | APC-Cy7 | RPA-T8 | 1 | BD | 557760 |
| Ttcr | 200 | CD45RA | BV421 | Hi100 | 1 | BD | 562885 |
| | | CD3 | Krome Orange | UCHT1 | 1 | BC | B00068 |
| | | TCRvd1 | FITC | TS8.2 | 1 | IV | TCR2730 |
| | | TCRvd2 | PE | B6 | 1 | BD | 555739 |
| | | TCRgd | PE-Cy5 | IMMU510 | 2 | BC | IM2662 |
| | | CCR7 | PC7 | 3D12 | 2 | BD | 560922 |
| | | TCRab | APC | IP26A | 1 | BC | B13981 |
| | | CD4 | Alexa700 | RPA-T4 | 1 | BD | 561030 |
| | | CD8a | APC-Cy7 | RPA-T8 | 1 | BD | 557760 |
| Tinv | 200 | CD16 | V450 | 3G8 | 2 | BD | 560475 |
| | | TCR Va7.2 | BV510 | 3C10 | 1 | BL | 351717 |
| | | CD3 | FITC | SK7 | 1 | BD | 345763 |
| | | iNKT | PE | 6B11 | 20 | BD | 552825 |
| | | CD8β | ECD | 2ST8.5H7 | 1 | BC | 6607123 |
| | | CD56 | PC7 | B159 | 1 | BD | 560916 |
| | | CD161 | APC | HP-3G10 | 1 | eB | 17-1619 |
| | | CD8α | Alexa700 | OKT8 | 1 | eB | 56-0086 |
| | | CD4 | APC-eFluor 780 | RPA-T4 | 1 | eB | 47-0049 |
| Ball | 200 | CD27 | V450 | M-T271 | 1 | BD | 560458 |
| | | CD19 | V500 | HIB19 | 2 | BD | 561121 |
| | | CD21 | FITC | BL13 | 1 | BC | IM0473U |
| | | CD10 | PE | ALB1 B9E9 | 10 | BC | A07760 |
| | | CD20 | ECD | (HRC20) | 1 | BC | IM3607U |
| | | CD38 | PC7 | HIT2 | 1 | eB | 25-0389 |
| | | CD24 | APC | ALB9 | 1 | BC | A87785 |
| | | IgD | Alexa700 | IA6-2 | 1 | BD | 561302 |
| Myel | 200 | CD16 | V450 | 3G8 | 2 | BD | 560475 |
| | | CD45 | V500 | 2D1 | 1 | BD | 655873 |
| | | CD33 | FITC | D3HL60.251 | 1 | BC | IM1135U |
| | | CD123 | PE | 9F5 | 1 | BC | B14808 |
| | | CD14 | ECD | RMO52 | 1 | BC | IM2707U |

| | | | | | | |
|---|---|---|---|---|---|---|
| | CD85k | PC5 | ZM3.8 | 1 | BC | IM3579 |
| | CD274 | PC7 | MIH1 | 2 | eB | 25-5983 |
| | CD86 | Alexa700 | 2331(FUN-1) | 1 | BD | 561124 |
| | HLA-DR | APC-H7 | G46-6 | 1 | BD | 561358 |

[1] BD, Becton Dickinson; BC, Beckman Coulter; bE, ebiosciences; BL, BioLegend ; IV, Invitrogen; RDS, R&D systems.

#### Cytometer and detectors

Cells were analyzed with a 10 color Gallios flow cytometer (Beckman-Coulter). The detectors used for each antibody specificity are described in the table.

**Table S1-3: Gallios' detectors and antigens corresponding to the eight staining conditions**

| Tube name | V450 | V550 | B525 | B575 | B620 | B695 | B770 | R660 | R725 | R770 |
|---|---|---|---|---|---|---|---|---|---|---|
| Trte | CD45RA | CD3 | CD31 | CD95 | CD27 | | CCR7 | CD28 | CD4 | CD62L |
| Tnai | CD45RA | CD3 | CD57 | CD279 | CD27 | | CCR7 | CD28 | CD4 | CD8α |
| Tact | CD45RA | CD3 | HLA-DR | CD127 | CD38 | | CCR7 | CD25 | CD4 | CD8α |
| Tpol | CD45RA | CD3 | CXCR5 | CCR6 | CCR4 | CD183 | CCR7 | | CD4 | CD8α |
| Ttcr | CD45RA | CD3 | TCRvδ1 | TCRvδ2 | | TCRγδ | CCR7 | TCRαβ | CD4 | CD8α |
| Tinv | CD16 | Vα7.2 | CD3 | iNKT | CD8b | | CD56 | CD161 | CD8α | CD4 |
| Ball | CD27 | CD19 | CD21 | CD10 | CD20 | | CD38 | CD24 | IgD | IgA |
| Myel | CD16 | | CD33 | CD123 | CD14 | CD85k | CD274 | | CD86 | HLA-DR |

#### Gating strategy

We used the Kaluza software (Beckman Coulter) for data analysis. Scatter signals are recorded as integral (INT, corresponding to peak area), peak (corresponding to peak height) and time of flight (TOF, corresponding to width).

##### *Step 1: Lymphocytes and myeloid cell definition*

We selected lymphocyte singlets by sequential gating strategy, as follows:

- A time/FSC INT gate defined events acquired at a constant flow rate;
- An SSC INT/FSC INT gate defined lymphocytes;
- An SSC INT/SSC PEAK gate defined singlet events;
- An FSC TOF/FSC INT gate further defined singlet events, by exclusion of residual doublets.

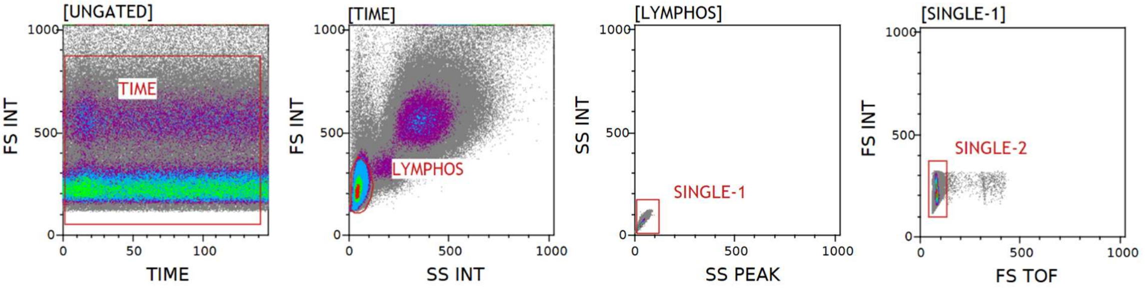

We selected myeloid cell singlets by sequential gating strategy, as follows:

- A time/FSC INT gate defined events acquired at a constant flow rate;
- An SSC INT/FSC INT gate defined lymphocytes and monocytes;
- An SSC INT/SSC PEAK gate defined singlet events;

- An FSC TOF/FSC INT gate further defined singlet events by exclusion of residual doublets.

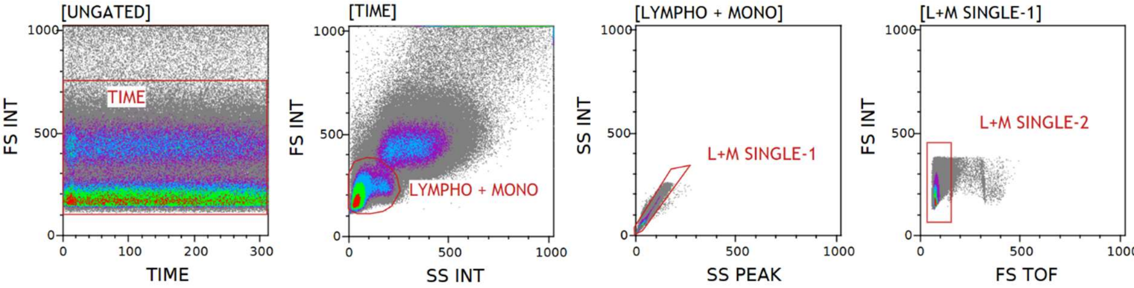

*Step 2: CD4 and CD8 T lymphocytes and their CD45RA/CCR7 subsets*

For the Trte, Tnai, Tact, Tpol and Ttcr tubes, we identified T lymphocyte subsets within lymphocyte singlets (SINGLE-2 gate, step 1). We used sequential gating strategy, as follows:

- A CD3/FSC INT gate defined T lymphocytes;

- CD4/CD8β gates defined CD4 ($CD4^{+}CD8\beta^{-}$) and CD8 ($CD4^{+}CD8\beta^{-}$ ) T lymphocytes;

- CD45RA/CCR7 quadrants defined four T lymphocyte subsets: naïve ($T_N$ , $CD45RA^{+}CCR7^{+}$), central memory ($T_{CM}$, $CD45RA^{-}CCR7^{+}$), effector memory ($T_{EM}$, $CD45RA^{-}CCR7^{-}$) and effector ($T_{EF}$, $CD45RA^{-}CCR7^{+}$).

- We defined CD4 and CD8 memory ($CD4T_M$ and $CD8T_M$) as non-naïve (non$CD45RA^{+}CCR7^{+}$).

We applied this classification for T lymphocyte labelled in Tact, Tpol and Ttcr panels. We used a more precise definition of T cell differentiation subsets for lymphocytes labelled in Tnai and Trte panels, as described in steps 3 and 4.

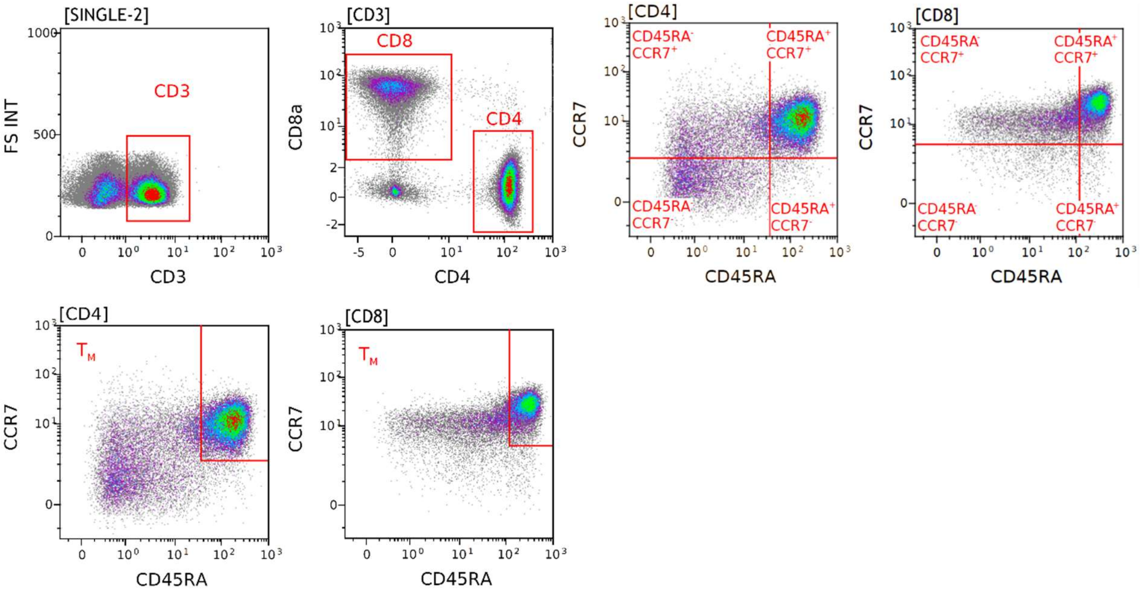

*Step 3: CD4 and CD8 T lymphocytes differentiation subsets defined by expression of CD45RA, CCR7, CD27 and CD28.*

We applied the gating strategy defined in steps 1 and 2. Then, we defined CD4 T lymphocyte subsets labeled in the Tnai tube, as follows:

- CD27/CD28 gates on CD45RA$^+$ (CD45RA$^+$R7$^+$ OR CD45RA$^+$R7$^-$) cells defined CD27$^+$CD28$^+$ naive T cells ($T_N$) and CD27$^-$CD28$^-$ effector T cells ($T_{EF}$);

- CD27/CD28 gates on CD45RA$^-$R7$^-$ cells defined CD27$^+$CD28$^+$ transitional memory T cells ($T_{TM}$) and CD27$^-$CD28$^{+/-}$ effector memory T cells ($T_{EM}$).

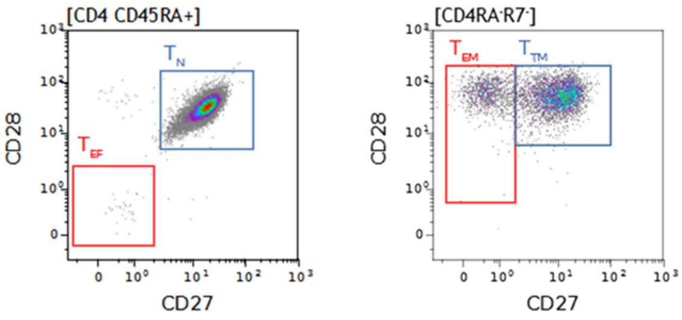

We further defined CD8 T lymphocyte subsets labeled in the Tnai tube by application of:

- CD27/CD28 gates on CD45RA$^+$ (CD45RA$^+$R7$^+$ OR CD45RA$^+$R7$^-$) cells defined CD27$^+$CD28$^+$ naive T cells ($T_N$) and CD27$^-$CD28$^-$ effector T cells ($T_{EF}$);

- CD27/CD28 quadrants defined four $T_{EM}$ subsets based on CD27 and CD28 expression.

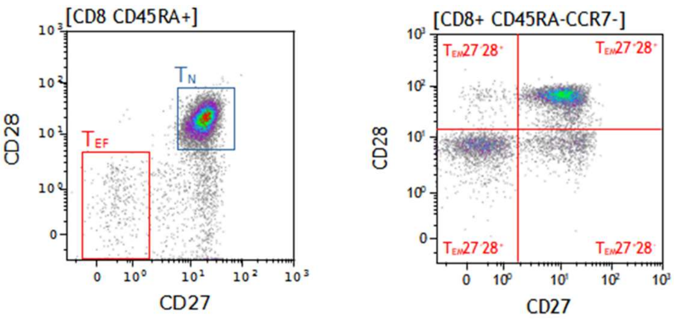

*Step 4: Recent thymic emigrants, stem cell memory and CD95+ memory CD4 T lymphocytes*

We applied the gating strategy defined in steps 1 and 2. Then, we defined CD4 T lymphocytes subsets labelled in the Trte tube, as follows:

- CD31/CD95 gates on CD45RA$^+$CCR7$^+$ cells defined CD31$^+$CD95$^-$ recent thymic emigrants ($T_{RTE}$), CD31$^-$CD95$^-$ naive T cells (CD31$^{neg}$$T_N$) and CD95$^+$ stem cell memory T lymphocytes ($T_{SCM}$);

- A CD95/CD62L gate on CD4 $T_M$ defined CD95$^+$ memory CD4 T lymphocytes (CD95$^+$$T_M$).

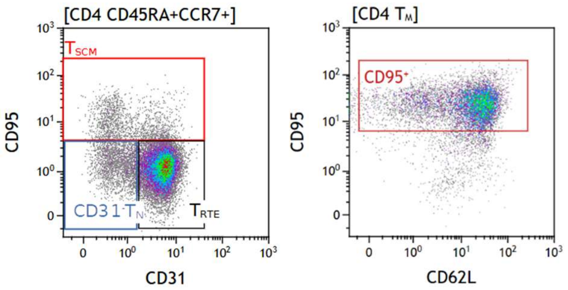

*Step 5: Activated CD4 and CD8 T lymphocytes defined by HLA-DR and CD38 coexpression*

We applied the gating strategy defined in steps 1 and 2. Then, we applied an HLA-DR/CD38 gate defining HLA-DR$^+$CD38$^+$ cells as activated memory CD4/CD8 T lymphocytes.

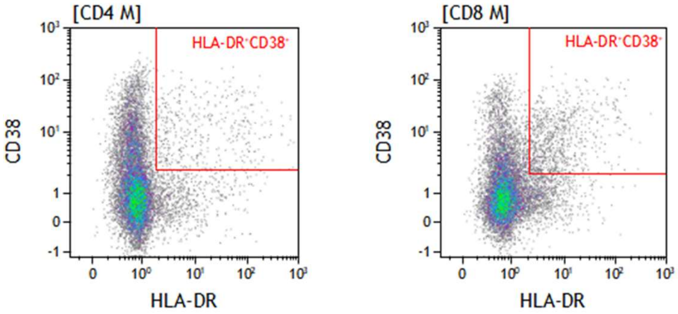

*Step 6. CD4 regulatory T lymphocytes defined by CD127 and CD25 markers and their differentiation and activated subsets*

We applied the gating strategy defined in steps 1 and 2. Then, we defined CD4 T lymphocytes labelled in the Tact tube by sequential gating strategy, as follows:

- A CD127/CD25 gate on CD4 T lymphocytes cells defined CD127$^-$CD25$^+$ cells as regulatory T lymphocytes ($T_{REG}$);
- Quadrants defined naïve, central memory, effector memory and effector CD4 $T_{REG}$ subsets based on CD45RA and CCR7 expression as described in step 2;
- We applied an HLA-DR/CD38 gate defining HLA-DR$^+$CD38$^+$ cells as activated CD4 $T_{REG}$.

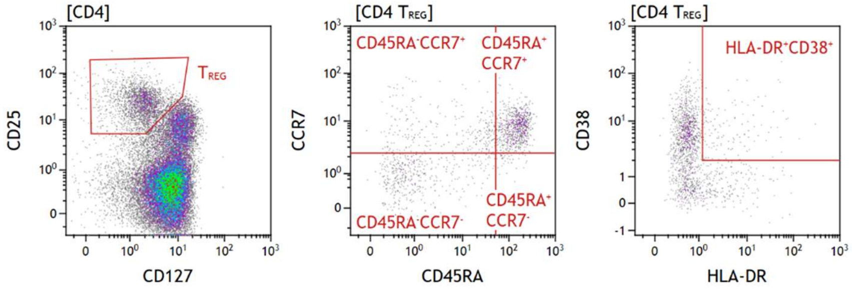

*Step 7. CD4 AND CD8 $T_{CM}$ and $T_{EM}$ lymphocytes defined by expression of chemokine receptors – The $CXCR5^+$ T lymphocytes*

We applied the gating strategy defined in steps 1 and 2. Then we characterized $T_{CM}$ and $T_{EM}$ lymphocytes. $T_N$ and $T_{EF}$ subsets rarely express chemokine receptors and were not included in the analysis.

Among $CXCR5^+$ CD4 T lymphocytes, we distinguished $CXCR3^+$ and $CXCR3^-$ populations using quadrant gates on CXCR5/CXCR3 dot plots. As $CXCR5^+$ CD4 T lymphocytes rarely expressed CCR4 or CCR6, we excluded these markers from further analysis.

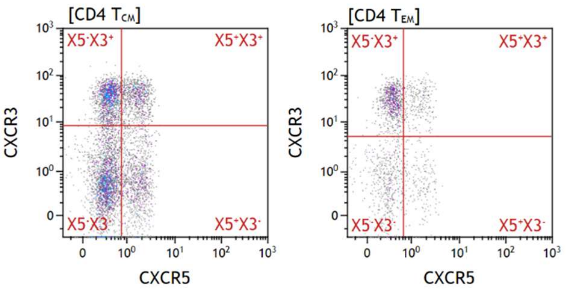

Among $CXCR5^+$ CD8 T lymphocytes, we distinguished $CCR4^+$ and $CCR4^-$ populations using quadrants gates on CXCR5/CCR4 dot plots. As $CXCR5^+$ CD8 T lymphocytes rarely expressed CXCR3 or CCR6, we excluded these markers from further analysis.

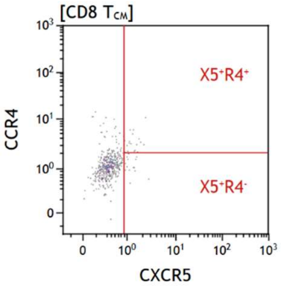

*Step 8. CD4 AND CD8 $T_{CM}$ and $T_{EM}$ lymphocytes defined by expression of chemokine receptors – The $CXCR5^-$ T lymphocytes*

For $CXCR5^-$ CD4 T lymphocytes, we applied CCR4/CCR6 quadrants on both $X5^-X3^+$ and $X5^-X3^-$ gates within CD4 $T_{CM}$ and CD4 $T_{EM}$ populations (step 7). We classified cells lacking CXCR3, CCR4, and CCR6 as non-polarized (Th0) helper cells. We classified $CXCR3^+$ cells as Th1, $CCR4^+$ cells as Th2, and $CCR6^+$ cells as Th17 cells, $CXCR3^+CCR4^+$ as Th1-2, $CXCR3^+CCR6^+$ Th1-17, $CCR4^+CCR6^+$ as Th2-17, and $CXCR3^+CCR4^+$ $CCR6^+$ as Th1-2-17 subsets.

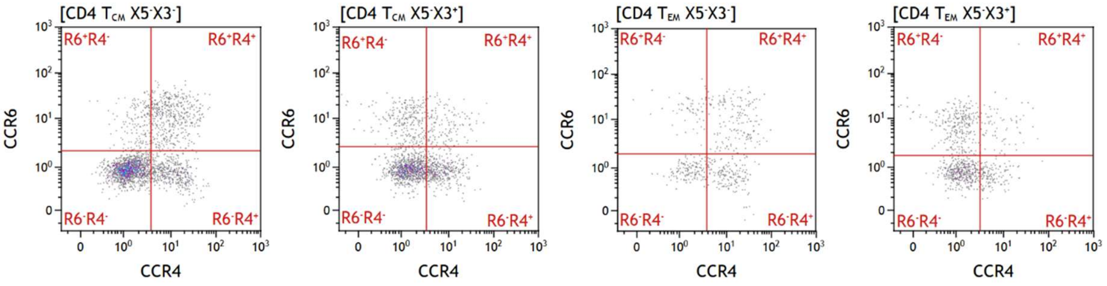

For CXCR5$^-$CCR4$^-$ CD8 $T_{EM}$ lymphocytes, we applied gates to define CXCR3 and CCR6 expression. We classified cells lacking CXCR3, CCR4, and CCR6 as non-polarized (Tc0) CD8 $T_{EM}$ lymphocytes. We classified CXCR3$^+$CCR6$^-$ cells as Tc1, CXCR3$^-$CCR6$^+$ cells as Tc17, and CXCR3$^+$CCR6$^+$ as Tc1-17 cells. As CXCR5$^-$ CD8 $T_{CM}$ lymphocytes rarely expressed CXCR3, CCR4 or CCR6, we excluded these markers from further analysis.

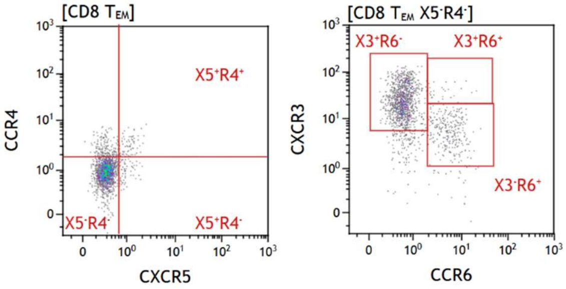

##### *Step 9. Gamma-delta T lymphocytes*

We selected T lymphocytes as CD3+ cells among lymphocyte singlets using the SINGLE-2 gate defined in step 1. We then gated TCRγδ$^+$ cells among CD3$^+$ lymphocytes. We did not analyze TCRαβ lymphocytes but used them as benchmark to position CD45RA/CCR7 quadrants (not shown). We defined γδ subsets using sequential gating strategy, as follows:

- Vδ1/Vδ2 gates identified TCR Vδ1$^+$, TCR Vδ2$^+$, and double negative TCRγδ;
- A CD8α/CD4 gate identified the CD8α$^+$ subset among TCR Vδ1$^+$ and TCR Vδ2$^+$;
- CD45RA/CCR7 quadrants identified naïve, central memory, effector memory and effector subsets (as defined in step 2) among TCR Vδ1$^+$ and TCR Vδ2$^+$.

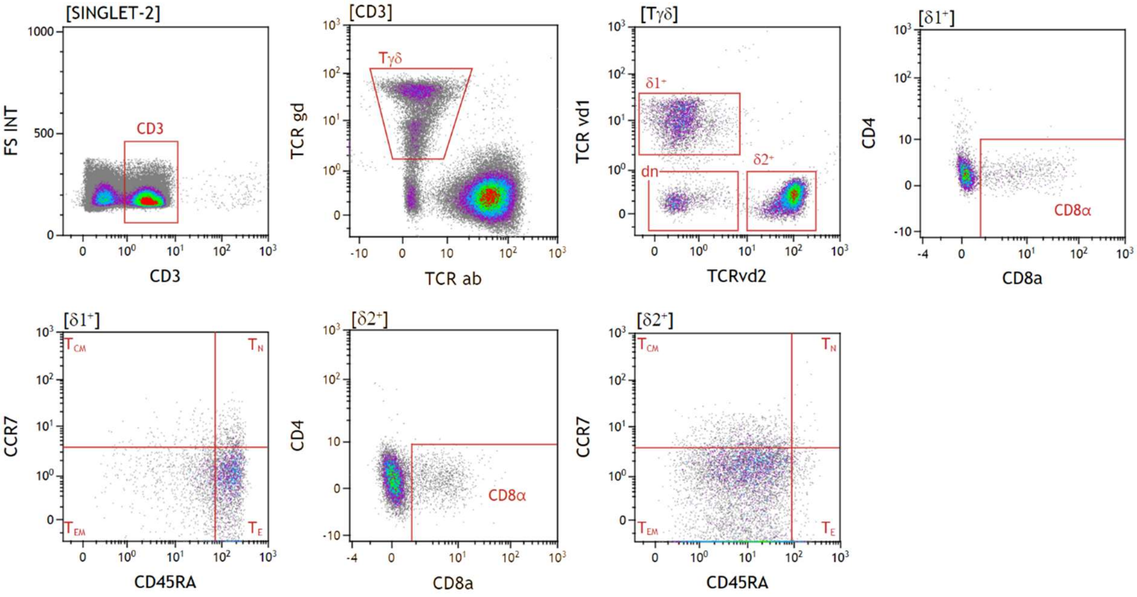

*Step 10. Unconventional T lymphocytes and NK lymphocytes*

Invariant natural killer T (iNKT) lymphocytes express the invariant TCRVα24-Jα18 T cell receptor (TCR). Mucosal-associated invariant T (MAIT) lymphocytes express the Vα7.2 segment joined with specific Jα segments (Jα12, Jα20, or Jα33) TCR [1]. We analyzed lymphocyte singlets using the SINGLE-2 gate defined in step 1, as follows:

- We defined iNKT lymphocytes as $CD3^{+}$ TCRVα24-$J\alpha18^{+}$ cells;
- We defined MAIT lymphocytes as CD3+ $TCRV\alpha7.2^{+}$ cells. Then, we sequentially applied CD8α/CD8β and CD161/CD4 gates to define the following five subsets: $CD161^{hi}CD8\alpha\beta$, $CD161^{-}CD8\alpha\beta$, $CD161^{hi}CD8\alpha\alpha$, $CD161^{hi}CD8^{-}$, and $CD161^{lo}CD4^{+}$;
- We selected $CD3^{-}$ lymphocytes and applied CD16/CD56 gates to define three natural killer (NK) lymphocyte subsets: $CD56^{HI}$, $CD56^{+}CD16^{+}$ and $CD56^{-}CD16^{+}$.

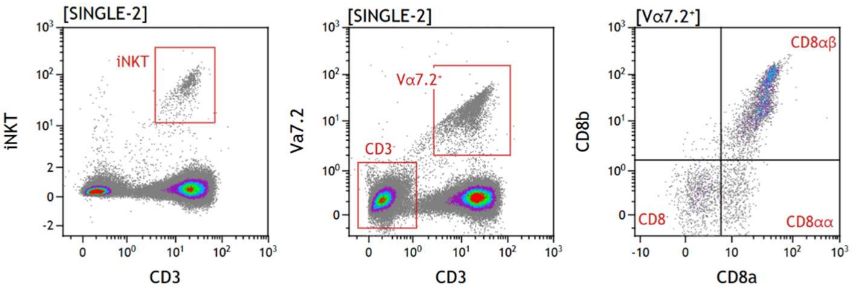

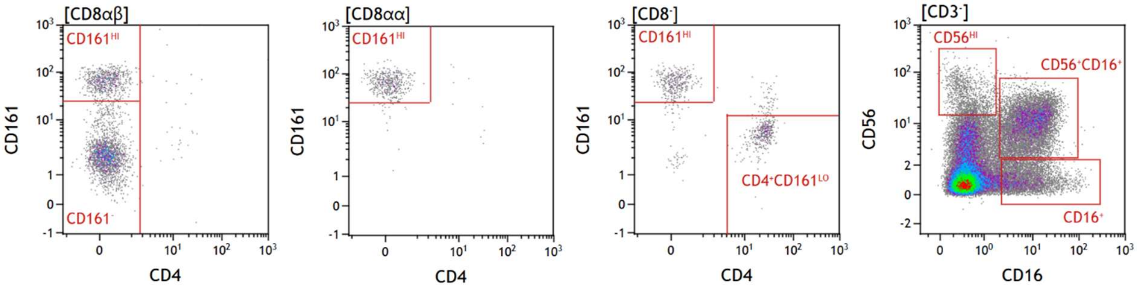

*Step 11. B lymphocytes*

We used the B lymphocyte classification from Moir and Fauci [2], except for the immature B lymphocytes for which we used the CD24 and CD38 markers [3].

We selected B lymphocytes as $CD19^+$ cells among lymphocyte singlets using the SINGLE-2 gate defined in step 1. We defined B cell subsets using sequential gating strategies, as follows:

- CD19/CD20 and CD10/CD27 gates identified $CD19^+CD20^+CD10^+CD27^+$ cells as resting memory B lymphocytes ($B_{RM}$);
- CD19/CD20, CD10/CD27 and CD24/CD38 gates identified $CD19^+CD20^+CD10^-CD27^-CD24^+CD38^+$ as immature B lymphocytes ($B_{IM}$);
- CD19/CD20, CD10/CD27 and CD21/CD38 gates identified $CD19^+CD20^+CD10^-CD27^-CD21^+CD38^+$ as naive B lymphocytes ($B_N$);
- Among $CD19^+CD20^+CD10^-CD27^+$ cells, the CD21/IgD quadrants identified $CD21^{low}IgD^-$ cells as exhausted tissue like memory B lymphocytes ($B_{ETLM}$), $CD21^+IgD^+$ cells as unswitched memory B cells ($B_{UM}$), and $CD21^+IgD^-$ cells as switched memory B cells ($B_{SM}$);
- Among $CD19^+CD20^+CD10^-CD27^-$ cells, the CD20/CD21 gate defined $CD20^+CD21^{low}$ tissue-like memory B lymphocytes ($B_{TLM}$);
- CD19/CD20 and CD27/CD38 gates identified $CD19^+CD20^-CD27^+CD38^{high}$ cells as plasmablasts ($B_{PB}$).

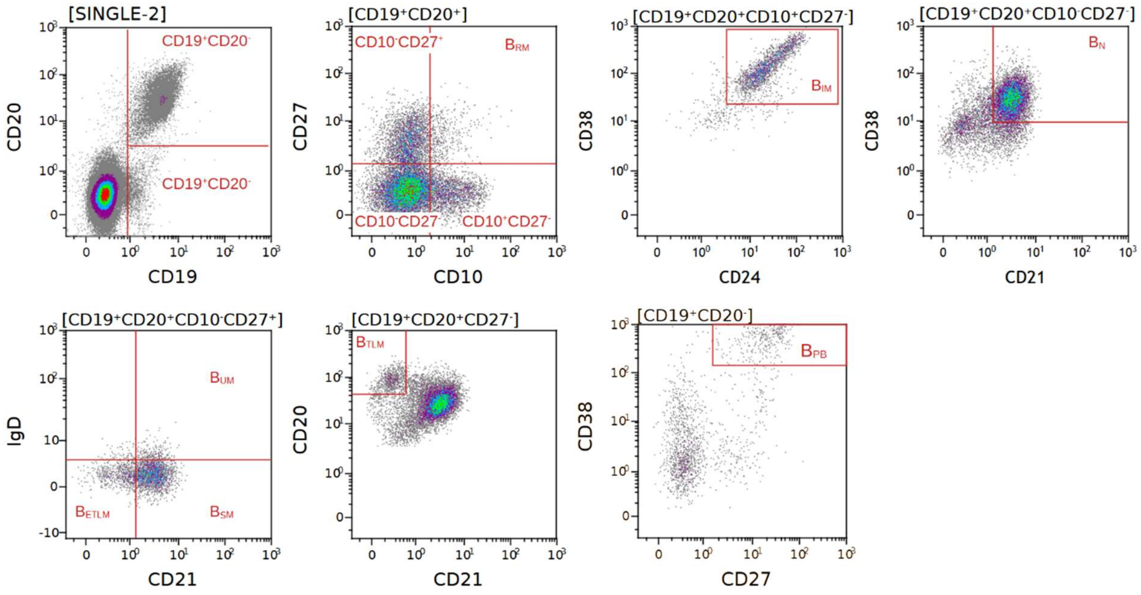

*Step 12. Monocytes and dendritic cells*

Myeloid cells, i.e. monocytes and dendritic cells, were selected among lymphocyte and monocyte singlets using the L+M SINGLE-2 gate defined on step 1.

Three CD14/CD16 gates defined classical $CD14^{+}CD16^{-}$ (MO1), intermediate $CD14^{+}CD16^{+}$ (MO2) and activated $CD14^{-/lo}CD16^{+}$ (MO3) monocyte subsets.

The sequential application of CD14/CD16 and HLA-DR/CD85j gates defined $CD14^{-}CD16^{-}HLA\text{-}DR^{+}CD85k^{+}$ cells as dendritic cells (DCs); CD33/CD123 gates defined $CD33^{+}CD123^{-}$ cells as myeloid DCs (mDCs) and $CD33^{-}CD123^{+}$ cells as plasmacytoid DCs (pDCs) [4,5].

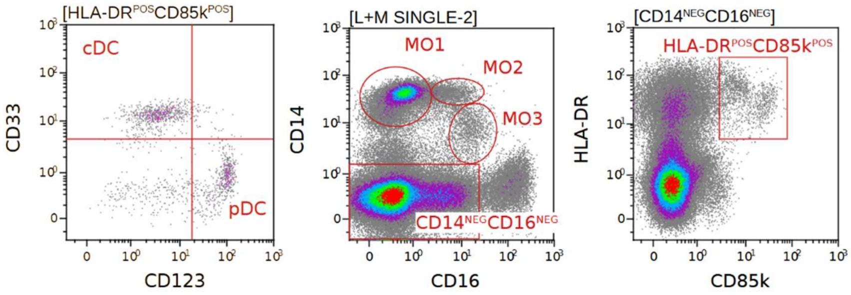

For each subset, we represented CD86, PD-L1 and HLA-DR expression levels on histograms and quantified them by the mean fluorescence intensity (MFI). Histograms are presented for the MO1 subset.

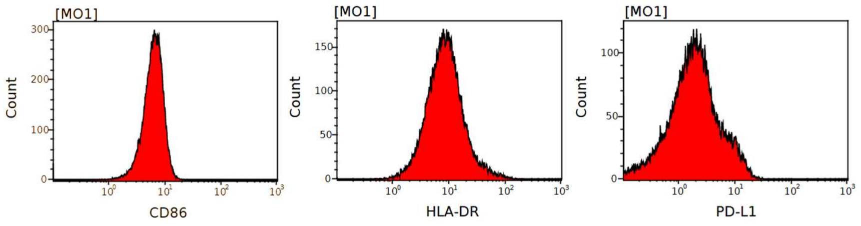

#### Memory PD-1$^+$CXCR5$^+$CXCR3$^-$ CD4 T lymphocytes detection using frozen PBMCs

We thawed frozen PBMCS and cultured them for 6 hours in DMEM supplemented with 10% human AB serum. We stained 5 x $10^5$ cells to quantify circulating PD-1$^+$CXCR5$^+$ memory CD4 $T_M$ lymphocytes, a resting memory population closely related to germinal center $T_{FH}$ cells [6]. The staining conditions and cytometer's detectors are summarized in tables S1-4 and S1-5, respectively. Cells were incubated for 30 min at RT in the dark, washed in PBS-0.1%BSA and resuspended in 0.4mL of PBS-2% paraformaldehyde. Data were immediately collected on a 13-color CytoFlex cytometer (Beckman Coulter).

**Table S1-4: staining conditions for the detection of CD4 $T_M$ expressing PD-1, CXCR5 and CXCR3**

| Antigen | Fluorochrome | Clone | Manufacturer | Catalog number | Quantity/tube (µL) |
|---|---|---|---|---|---|
| Live-dead | Yellow | | MP | L34967 | 0.5 |
| CD3 | Krome Orange | UCHT1 | BC | B00068 | 5 |
| CD4 | AF 700 | RPA-T4 | BD | 557922 | 1 |
| CD45RA | BV421 | HI100 | BD | 562885 | 1 |
| CCR7 | PC7 | 3D12 | BD | 557648 | 2 |
| CD279 (PD-1) | PE | J105 | eB | 12-2799-41 | 1 |
| CD183 | PC5 | 1C6 | BD | 561731 | 2 |
| CXCR5 | FITC | 51505 | RDS | FAB190F | 5 |

[1] BD, Becton Dickinson; BC, Beckman Coulter; eB, ebiosciences; MP, molecular probes; RDS, R&D systems.

**Table S1-5: Cytoflex' detectors and antigens used for the detection of CD4 $T_M$ expressing PD-1, CXCR5 and CXCR3**

| Detector | Antigen |
|---|---|
| V660 | |
| V610 | Live-dead |
| V525 | CD3 |
| V450 | CD45RA |
| B690 | |
| B525 | CXCR5 |
| Y780 | CCR7 |
| Y690 | CD183 |
| Y610 | |
| Y585 | CD279 |
| R780 | |
| R712 | CD4 |
| R660 | |

We acquired data on a 13 color CytoFlex cytometer and analyzed them with the Kaluza software (Beckman Coulter). We used the Kaluza software (Beckman Coulter) for data analysis. Scatter signals are recorded as integral (INT, corresponding to peak area) and peak (corresponding to peak height). The gating strategy sequentially defined viable lymphocytes (see above), memory CD4 T lymphocytes, and PD-1$^+$CXCR5$^+$CXCR3$^-$ cells within this population.

Step 1: We selected single viable lymphocytes by sequential gating strategy, as follows:

- A time/FSC INT gate defined events acquired at a constant flow rate;

- An SSC INT/FSC INT gate defined lymphocytes;

- A live-dead/FSC INT gate defined viable lymphocytes;

- An SSC INT/SSC PEAK gate defined single viable lymphocytes.

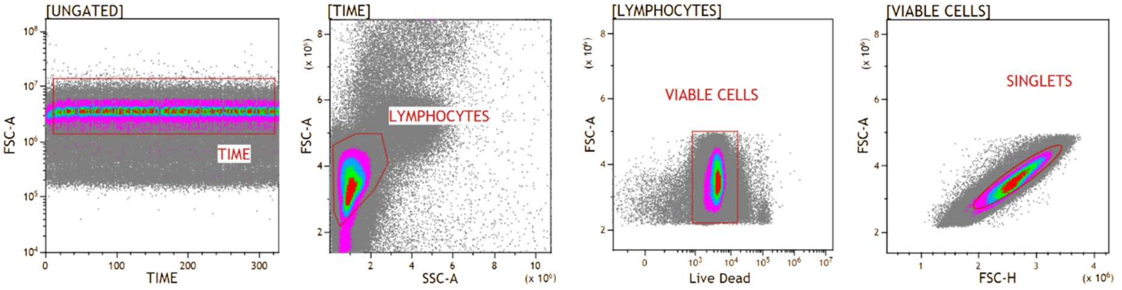

Step 2. We selected PD-1$^+$CXCR5$^+$ memory CD4 $T_M$ lymphocytes by sequential gating strategy, as follows:

- A CD3/CD4 gate on singlet cells defined CD4+ T lymphocytes;

- A CD45RA/CCR7 gate defined memory (CD45RA$^-$) CD4 T lymphocytes;

- A PD-1/CXCR5 gate defined PD-1$^+$CXCR5$^+$ memory CD4 T lymphocytes;

- A CXCR3/CXCR5 gate defined memory PD-1$^+$CXCR5$^+$CXCR3$^-$ CD4 T lymphocytes.

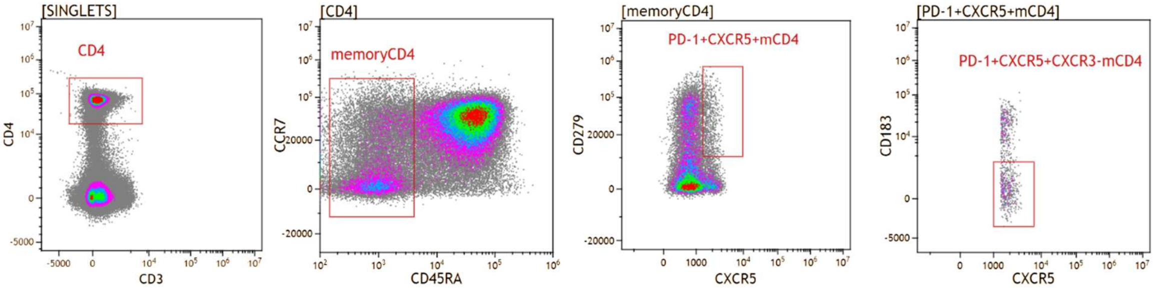

#### *In vitro* memory B cell stimulation and anti-HIV IgG production.

##### PBMCs culture

We used frozen PBMCs for this assay, cultured in RPMI supplemented with 10% fetal calf serum and 10 mM HEPES. We added DNase I to prevent cell clumping (5 U/mL, Invitrogen 18047-019). After a resting period of 6 hours, PBMCS were enumerated, seeded at $10^5$ cells / well in U-bottom 96-well plate and stimulated with R848 (500 ng/mL, Miltenyi 130-109-376) and IL-2 (100U/mL,Miltenyi 130-097-744). IL-6 (10 ng/mL, Miltenyi 130-095-352) was added in half of stimulated wells. Seven days post-stimulation, cells from 3 wells were collected and stained for the identification of CD19$^+$B cells and CD27$^{hi}$CD38$^{hi}$CD21$^{lo}$CD19$^+$ plasmablasts (see below).

#### Total and HIV-specific IgG quantification

We quantified secreted IgG levels in culture supernatants by ELISA. We coated clear high-binding flat-bottom polystyrene 96-well microplates (R&D Systems, DY990) overnight at 4°C with monoclonal rabbit anti-human Ig light chain antibody (0.5 µg/mL, Sigma-Aldrich, SAB5600122). We blocked free binding sites with PBS–0.1% Tween 20–1% BSA for 1h30 at RT. We diluted supernatants in carbonate buffer (1:10, 1:100, and 1:1000) and created a titration curve ranging from 10 to 0.0024 µg/mL using purified IgG from human serum (Sigma-Aldrich, I4506). We used wells coated with carbonate buffer to generate blank values. We incubated samples and the standard curve for 1 h at RT, then added horseradish peroxidase-conjugated goat anti-human IgG HL (1:40,000; Jackson Immuno Research Europe, 109-035-008) for 1 h at RT, followed by 0.25 mg/mL o-phenylenediamine dihydrochloride in 0.05 M citrate buffer (pH 5) and 0.012% $H_2O_2$ for 30 min. We performed all incubations in a humidified chamber and washed plates with PBS-0.05% Tween 20 between steps. We stopped the reaction by adding 1.8 M $H_2SO_4$ (vol/vol) and read absorbance at 493 nm. We tested all samples in duplicate and used average optical density (OD) values. We subtracted blank well values from standard and test wells. We calculated a four-parameter sigmoid curve of corrected OD as a function of IgG concentration using Prism software (version 10, GraphPad Software) and used this curve to determine IgG concentrations in supernatants. We detected HIV-specific IgG in an ELISA assay using the protocol described above for IgG quantification, with the following modifications. We coated ELISA plates with recombinant HIV proteins (1 µg/mL HIV-1 IIIB p24, #12028; 0.5 µg/mL HIV-1 UG037 gp140, #12063; 0.5 µg/mL HIV-1 IIIB gp120, #12027; NIH AIDS Reagent Program) diluted in carbonate buffer. We diluted culture supernatants 1:2 in a 1:1 mixture of PBS and low-cross buffer (Candor, 100-035). We generated a titration curve using a mixture of two plasma samples from HIV-1-infected adolescents [7] (starting at a 1:100 plasma dilution with serial 3-fold dilutions). The corrected OD value obtained with the 1:100 diluted plasma mixture corresponds to 1 arbitrary unit.

#### B lymphocyte and plasmablast detection at day 7 post-stimulation

B lymphocytes and plasmablast expansion was assessed by flow cytometry and we defined plasmabalsts as $CD27^{hi}CD38^{hi}CD21^{lo}$ B lymphocytes [2]. Seven days post-stimulation, cells from 3 culture wells were collected in 5 mL tubes, washed in PBS-0.1%BSA and stained for the identification of B lymphocytes and plasmablasts. The staining conditions and cytometer's detectors are summarized in tables S1-6 and S1-7, respectively. Cells were incubated for 30 min at RT in the dark, washed in PBS-0.1%BSA and resuspended in 0.4mL of PBS-2% paraformaldehyde. Data were immediately collected on a 13 color CytoFlex cytometer (Beckman Coulter).

**Table S1-6: staining conditions for the detection of B lymphocytes and plasmablasts.**

| Antigen | Fluorochrome | Clone | Quantity/tube (µL) | Manufacturer | Catalog number |
|---|---|---|---|---|---|

| | | | | | |
|---|---|---|---|---|---|
| live-dead | green | | 1 | MP | L23101 |
| CD3 | PE | UCHT1 | 2 | BD | 561808 |
| CD19 | V500 | HIB19 | 2 | BD | 561121 |
| CD21 | BB700 | B-ly4 | 2 | BD | 566569 |
| CD27 | V450 | M-T271 | 1 | BD | 560458 |
| CD38 | PC7 | HIT2 | 1 | eB | 25-0389 |

[1] BD, Becton Dickinson; eB, ebiosciences; MP, molecular probes.

**Table S1-7: Cytoflex' detectors and antigens used for the detection of CD4 $T_M$ expressing PD-1, CXCR5 and CXCR3**

| Detector | Antigen |
|---|---|
| V660 | |
| V610 | |
| V525 | CD19 |
| V450 | CD27 |
| B690 | CD21 |
| B525 | live-dead |
| Y780 | CD38 |
| Y690 | CD3 |
| Y610 | |
| Y585 | |
| R780 | |
| R712 | |
| R660 | |

We analyzed data with the Kaluza software (Beckman Coulter). Scatter signals are recorded as integral (INT, corresponding to peak area), and peak (corresponding to peak height). The gating strategy sequentially defined viable single lymphocytes and lymphoblastes, CD19+ B lymphocytes, and $CD21^{lo}CD27^{hi}CD38^{hi}$ plasmablasts.

Step 1: We selected single viable lymphocytes by sequential gating strategy, as follows:

- A time/FSC INT gate defined events acquired at a constant flow rate;
- An SSC INT/FSC INT gate defined lymphocytes;
- A live-dead/FSC INT gate defined viable lymphocytes;
- An SSC INT/SSC PEAK gate defined single viable lymphocytes.

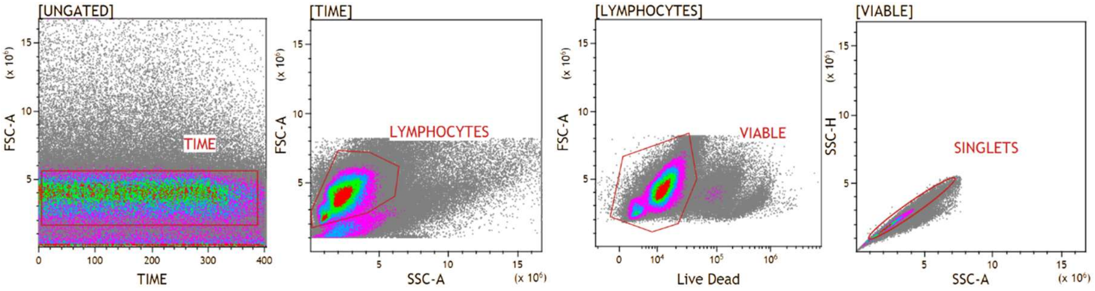

STEP 2: We selected B lymphocytes and plasmablasts by sequential gating strategy, as follows:

- A CD3/CD19 gate applied to single lymphocytes defined $CD19^{+}$ B lymphocytes;

- A CD27/CD38 gate defined CD27$^{hi}$CD38$^{hi}$ B lymphocytes

- A CD21/CD27 gate defined CD27$^{hi}$CD38$^{hi}$CD21$^{lo}$ cells as plasmablasts.

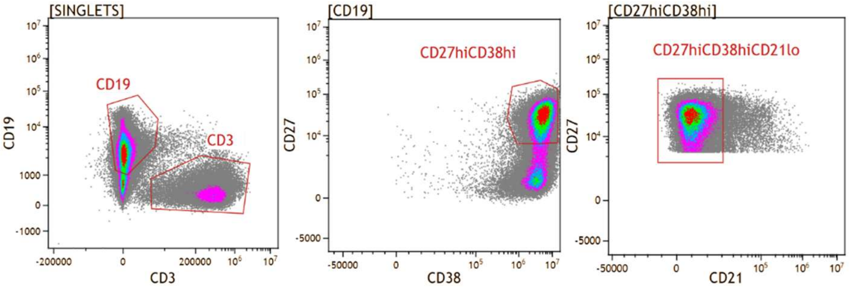

#### Phenotypic study of natural killer lymphocytes performed on fresh blood samples

##### Staining conditions

We performed natural killer (NK) lymphocyte phenotyping on fresh blood samples collected on EDTA tubes for 63 out of the 76 participants. We stained 100 µL of whole blood with six mixtures of antibodies in 5 mL tubes (Becton Dickinson, cat. no. 352054) for 30 min at RT in the dark. We then added 3 mL of red blood cell lysis kit (eBioscience, cat. no. 00-5333-57), mixed samples by low speed vortexing and incubated the cells 15 min at RT. The staining conditions and cytometer's detectors are summarized in tables S1-8 and S1-9, respectively. We centrifuged the samples for 6 min at 500 x g, washed the pellet in 2 mL PBS, resuspended in 0.3 mL PBS and immediately analyzed. The table describes the six sets of staining conditions.

**Table S1-8: the six sets of staining conditions for NK lymphocyte analyses**

| Tube name | Blood volume, µL | Antigen | Fluorochrome | Clone | Volume, µl | Manufacturer[1] | Cat N° |
|---|---|---|---|---|---|---|---|
| NK1 to NK6 | 100 | CD16 | Pacific blue | 3G8 | 3 | BC | B36292 |
| | | CD3 | ECD | UCHT1 | 2 | BC | A07748 |
| | | CD56 | Alexa700 | B159 | 3 | BD | 557919 |
| | | CD45 | Krome Orange | J33 | 3 | BC | B36294 |
| NK1 | | CD69 | FITC | FN50 | 15 | BD | 555530 |
| | | CD160 | PE | BY55 | 5 | BC | IM3657 |
| | | CD159c (NKG2C) | APC | 134591 | 4 | RDS | FAB138A |
| NK2 | | CD94 | FITC | HP-3D9 | 3 | BD | 555888 |
| | | CD85j (ILT2) | PE | HP-F1 | 5 | BC | A07408 |
| | | CD314 (NKG2D) | APC | BAT221 | 4 | MB | 130-099-215 |
| NK3 | | CD158d (KIR2DL4) | FITC | 181703 | 3 | RDS | FAB2238F |
| | | CD337 (NKp30) | PE | Z25 | 5 | BC | IM3709 |
| | | CD335 (NKp46) | APC | 195314 | 4 | RDS | FAB1850A |
| NK4 | | CD226 (DNAM) | FITC | DX11 | 3 | BD | 559788 |
| | | CD161 | PC5 | DX12 | 2 | BD | 551138 |
| | | CD336 (NKp44) | PE | Z231 | 5 | BC | IM3710 |
| | | NKp80 | APC | 239127 | 4 | RDS | FAB1900A |

| | | | | | | |
|---|---|---|---|---|---|---|
| NK5 | CD158e1 (KIR3DL1) | FITC | DX9 | 3 | BD | 555966 |
| | CD158e1/e2 (KIR3DL1/DS1) | PE | Z27.3.7 | 5 | BC | IM3292 |
| | CD159a (NKG2A) | APC | 131411 | 4 | RDS | FAB1059A |
| NK6 | CD158a (KIR2DL1) | FITC | HP-3E4 | 3 | BD | 556062 |
| | CD244 (2B4) | PC5 | C1.7 | 3 | BC | IM2658 |
| | CD158i (KIR2DS4) | PE | FES172 | 5 | BC | IM3337 |
| | CD158b (KIR2DL2/3) | APC | GL183 | 4 | BC | A22333 |

[1] BD, Becton Dickinson; BC, Beckman Coulter; MB, Miltenyi Biotec; RDS, R&D systems.

#### Cytometer and detectors

Cells were analyzed with a 10 color Gallios flow cytometer (Beckman-Coulter). The detectors used for each antibody specificity are described in the table.

**Table S1-9: Detectors and antigens corresponding to the six staining conditions**

| Tube name | V450 | V550 | B525 | B575 | B620 | B695 | B770 | R660 | R725 | R770 |
|---|---|---|---|---|---|---|---|---|---|---|
| NK1 | CD16 | CD45 | CD69 | CD160 | CD3 | | | CD159c | CD56 | |
| NK2 | CD16 | CD45 | CD94 | CD85j | CD3 | | | CD314 | CD56 | |
| NK3 | CD16 | CD45 | CD158d | CD337 | CD3 | | | CD335 | CD56 | |
| NK4 | CD16 | CD45 | CD226 | CD336 | CD3 | CD161 | | NKp80 | CD56 | |
| NK5 | CD16 | CD45 | CD158e1 | CD158e1/e2 | CD3 | | | CD159a | CD56 | |
| NK6 | CD16 | CD45 | CD158a | CD158I | CD3 | CD244 | | CD158b | CD56 | |

#### Gating strategy

We used the Kaluza software (Beckman Coulter) for data analysis. Scatter signals are recorded as integral (INT, corresponding to peak area).

##### *Step 1: Natural killer lymphocytes definition*

We identified NK lymphocytes using sequential gating strategy, as follows:

- A CD45/FSC INT gate defined $CD45^+$ cells;
- An SSC INT/FSC INT gate defined lymphocytes;
- A CD3/FSC INT gate defined $CD3^-$ lymphocytes;
- CD16/CD56 gates defined five NK lymphocyte subsets.

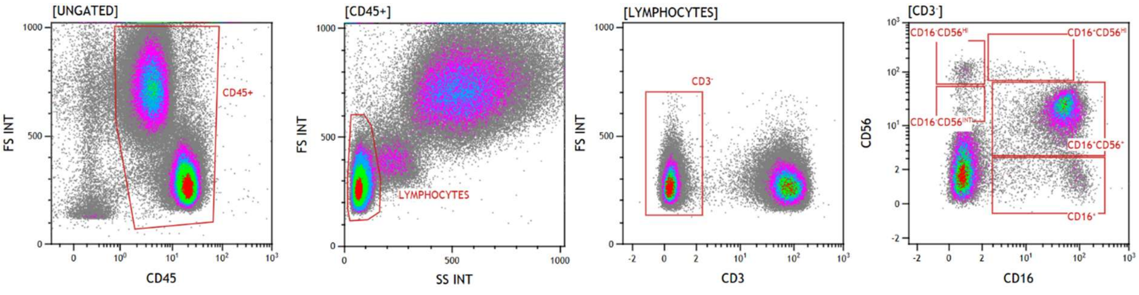

##### *Step 2. Natural killer lymphocyte receptors*

We defined total NK lymphocytes as a boolean gate encompassing five subsets: $CD16^-CD56^{HI}$ OR $CD16^-CD56^{INT}$ OR $CD16^+CD56^{HI}$ OR $CD16^+CD56^+$ OR $CD16^+$. We quantified NK receptor expression as the percentage of positive cells on histograms except for CD85j and NKG2A for which we defined subsets expressing low or high levels of these molecules. For receptors lacking a clear bimodal or trimodal distribution, we used dot plots as a reference to determine the positivity threshold (the CD69/CD160 dot plot from tube 1 is presented as an example). We used two different antibodies to stain the CD158e molecules: the Z27.3.7 antibody recognizes both CD158e1 (KIR2DL1) and CD158e2 (KIR2DS1) while the DX9 antibody recognizes CD158e1 only.

##### NK receptors – Tube 1

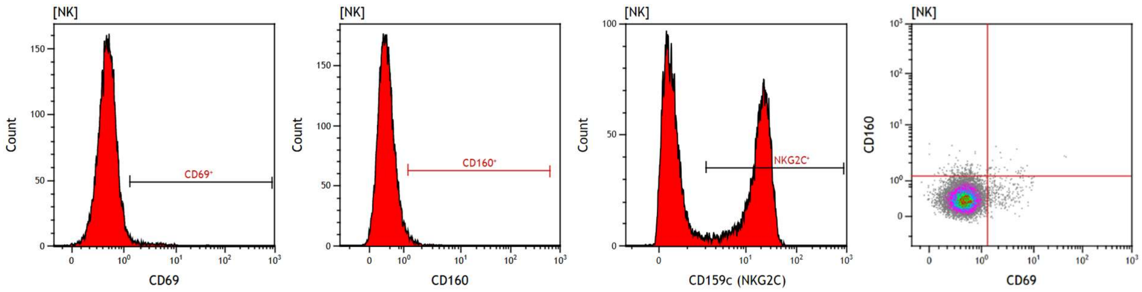

##### NK receptors – Tube 2

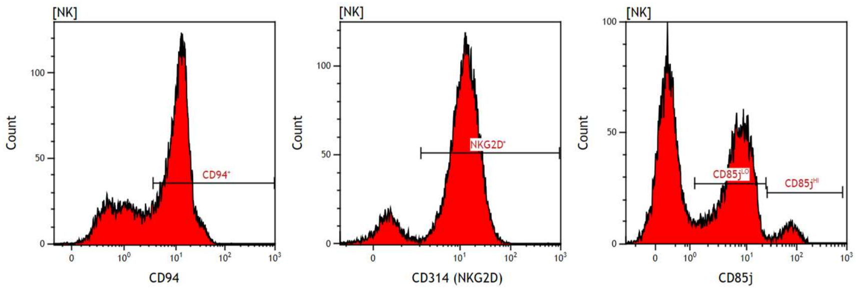

##### NK receptors – Tube 3

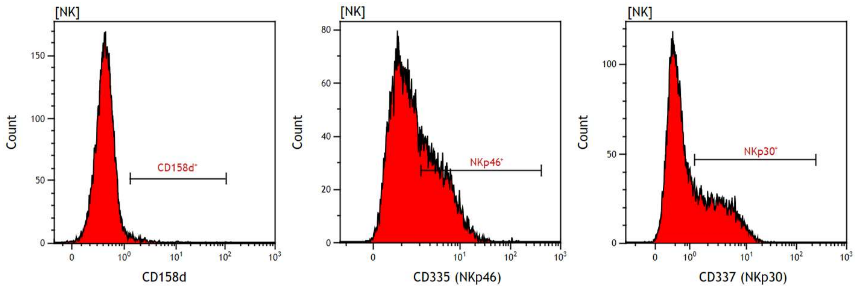

NK receptors – Tube 4

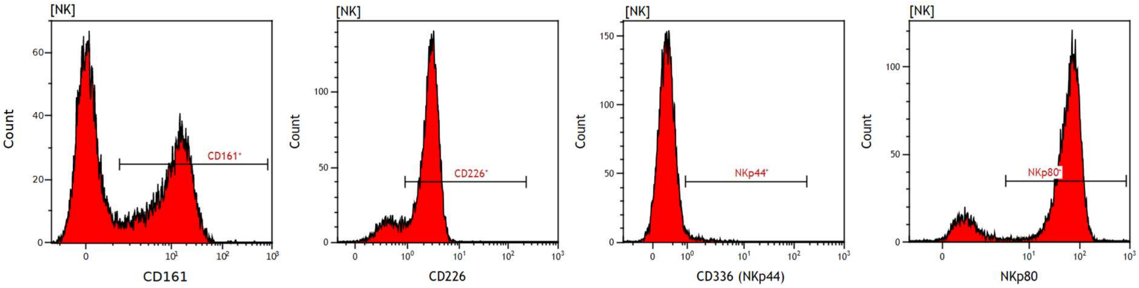

NK receptors – Tube 5

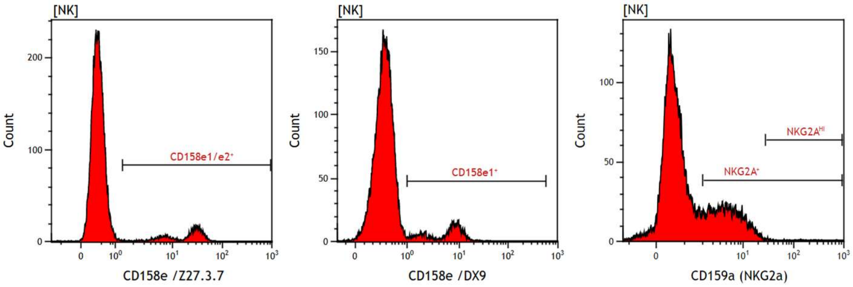

NK receptors – Tube 6

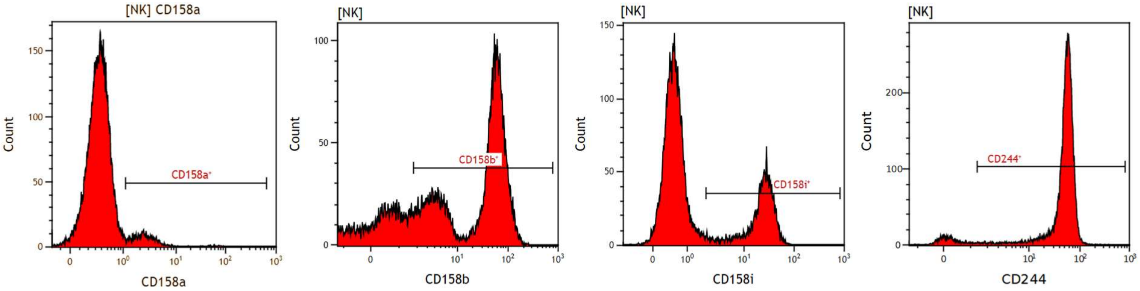

### T-lymphocyte functional assays

#### Material and methods

##### *ELISpot*

PBMCs were isolated from whole heparinized blood by density gradient centrifugation over Ficoll-Paque (Eurobio-Scientific, CMSMSL01-01). The ELISpot for analysis of IFN-γ production was carried out on fresh PBMCs. Multiscreen IP 96-well plates with PVDF membrane (MSIPS4510; Millipore) were pre-wetted with 35% ethanol for 1 min at RT, washed with PBS and coated overnight at 4°C with 50 µL of anti-IFN-γ capture antibody from the Diaclone kit # 869.060.010 in PBS. The antibody-coated plates were then washed three times with PBS, blocked with PBS containing 5% fetal calf serum for at least 2h at RT, and washed three times with PBS. PBMCs resuspended in RPMI 1640 (Gibco, 61870010) supplemented with 10% fetal calf serum (Biowest). Cell densities were seeded at 2 x $10^5$ and 5 x $10^4$

cells/well for peptide stimulation, or 5 x $10^4$ and 1.25 x $10^4$ cells/well for mitogen stimulation. HIV-1 potential T-cell epitope (PTE) Gag and Nef peptide pools (NIH AIDS reagent program, #12437 and #12822) and HCMV pp65 peptide pool (#11549) were added at a final concentration of 1 µg of each peptide/mL. Negative controls consisted of cells incubated in medium and peptide diluent (0.2% DMSO, Sigma-Aldrich D4540); positive controls were cells stimulated with phorbol myristate acetate (PMA, 25 ng/mL, Sigma-Aldrich Chimie, P8139) and ionomycin (100 ng/mL, Sigma-Aldrich Chimie, I3909). The cells were incubated overnight at 37°C with 5% $CO_2$. Wells were washed three times in PBS with 0.05% Tween 20 and three times with PBS and incubated 3-4 hour at RT with 50µL of the biotinylated anti-IFN-γ detection antibody in PBS. After washing as described above, streptavidin-conjugated alkaline phosphatase (1µg/mL, Amersham, AMDEX RPN4402) diluted in PBS – 1%BSA was added for 90 min at RT. The wells were washed, and 5-bromo-4-chloro-3-indolylphosphate–Nitro Blue Tetrazolium color substrate (BCIP/NBT, Promega, S3771) diluted in 100 mM Tris, 150 mM NaCl, and 5 mM MgCl2 (pH 9.5) was added for 30 min at room temperature before the reaction was stopped with tap water. IFN-γ spot forming cells (SFCs) were enumerated using an S6 Ultimate Image UV analyzer (CTL Europe, Bonn, Germany). A minimum of 10 spots/well were retained to calculate the frequencies of IFN-γ SFCs, corresponding to a detection threshold of 50 IFN-γ SFCs /$10^6$ PBMCs.

*Intracellular cytokine staining*

PBMCs were thawed and cultured in RPMI supplemented with 10% fetal calf serum and 10 mM HEPES. DNAse I (5U/mL, Invitrogen 18047-019) was added to prevent cell clumping. After 6-hour resting period, PBMCS were enumerated, seeded at 4 x $10^5$ cells/well in conical bottom 96-well plates and stimulated with HIV-1 potential T-cell epitope (PTE) Gag and Nef peptide pools (NIH AIDS reagent program, #12437 and #12822) and HCMV pp65 peptide pool (#11549) at 1 µg/mL of each peptide Negative controls consisted of cells incubated with medium and peptide diluent (0.2% DMSO, Sigma-Aldrich D4540); positive controls consisted of cells stimulated with Dynabeads® Human T-Activator CD3/CD28 (5µL/well, Gibco, 11132D). Brefeldin A (4µg/mL, Biolegend, #460201) was added to block cytokine secretion, and cells were incubated overnight at 37°C with 5% $CO_2$. Sixteen hours post-stimulation, cells were washed, resuspended in PBS - 0.1% BSA and stained with antibodies targeting the CD3, CD4, CD8, CD154 (CD40L) molecules and a viability marker for 15 min at RT (Table). Cells were then fixed and permeabilized using the Fix & Perm kit (Invitrogen, GAS003) and stained with antibodies targeting IL-2, TNF-α, MIP-1β and IFN-γ for 15 min at RT. The staining conditions and cytometer's detectors are summarized in tables S1-10 and S1-11, respectively. Cells were washed in PBS-BSA 0.1% and resuspended in PBS – 0.1% paraformaldehyde and data were immediately collected on a 13 color CytoFlex cytometer (Beckman Coulter). As detailed below, the gating strategy defined CD4 and CD8 T lymphocytes expressing each of the four cytokines. A positivity threshold of 0.1% cytokine-positive cells was applied, based on background responses observed in uninfected donors.

**Table S1-10: staining conditions for the detection of cytokine production by T lymphocytes**

| Antigen | Fluorochrome | Clone | Quantity/tube (µL) | Manufacturer[1] | Catalog number |
|---|---|---|---|---|---|
| Surface staining | | | | | |
| live-dead | Aqua | | 0.5 | MP | L34957 |
| CD3 | PE594 | UCHT1 | 2 | BL | 300450 |
| CD4 | Alexa Fluor700 | RPA-T4 | 1 | BD | 557922 |
| CD8 | APC-Cy7 | RPA-T8 | 1 | BD | 557760 |
| CD40L | BB700 | TRAP1 | 2.5 | BD | 745814 |
| Intracellular staining | | | | | |
| IL-2 | BV605 | MQ1-17H12 | 5 | BD | 564165 |
| TNF-α | FITC | MAb11 | 1 | BD | 554512 |
| MIP1-β | PE | D21-1351 | 1 | BD | 550078 |
| IFN-γ | APC | B27 | 1 | BD | 554702 |

[1] BD, Becton Dickinson; BC, Beckman Coulter; bE, ebiosciences; BL, BioLegend ; IV, Invitrogen; MP, molecular probes; RDS, R&D systems.

**Table S1-11: Cytoflex' detectors and antigens used for the detection of CD4 $T_M$ expressing PD-1, CXCR5 and CXCR3**

| Detector | Antigen |
|---|---|
| V660 | |
| V610 | live-dead |
| V525 | CD3 |
| V450 | CD45RA |
| B690 | |
| B525 | CXCR5 |
| Y780 | CCR7 |
| Y690 | CD183 |
| Y610 | |
| Y585 | CD279 |
| R780 | |
| R712 | CD4 |
| R660 | |

#### Gating strategy

We used the Kaluza software (Beckman Coulter) for data analysis. Scatter signals are recorded as integral (INT, corresponding to peak area), and peak (corresponding to peak height).

Step 1: We selected single viable lymphocytes by sequential gating strategy, as follows:

- A time/FSC INT gate defined events acquired at a constant flow rate;

- An SSC INT/FSC INT gate defined lymphocytes;

- A live-dead/FSC INT gate defined viable lymphocytes;

- An SSC INT/SSC PEAK gate defined single viable lymphocytes.

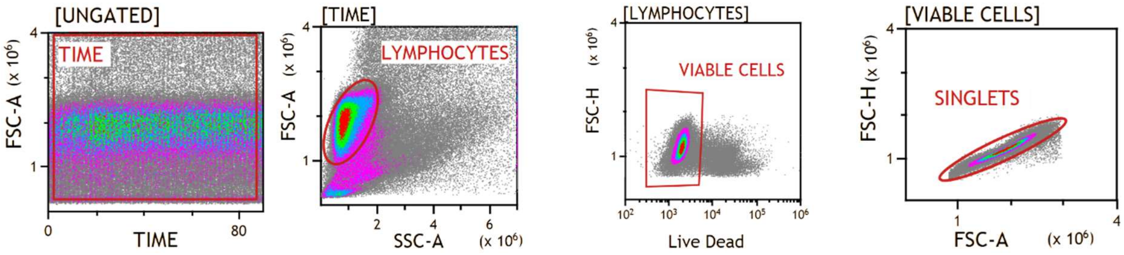

Step 2: CD4 and CD8 T lymphocytes expressing CD40L, IL-2, TNF-α, MIP-1β or IFN-γ were selected, as follows:

- A FSC INT/CD3 gate applied to events selected by SINGLETS gate (step 1) defined $CD3^+$ T lymphocytes;
- Two CD4/CD8 gates defined $CD4^+CD8^-$ and $CD4^-CD8^+$ T lymphocytes;
- A CD40L/CD8 gate applied to CD3+ T lymphocytes to define $CD8^-$ (i.e., CD4) T lymphocytes expressing CD40L;
- IL-2/CD8, TNF-α/CD8, MIP-1β/CD8, and IFN-γ/CD8 gates each defined $CD4^+$ ($CD8^-$) and $CD8^+$ T cells expressing the respective cytokine.

CD40L coexpression with each cytokine was used to precisely identify antigen-responsive $CD4^+$ T cells. For each cytokine, the corresponding CD40L-CD4 and cytokine-CD4 gates were combined using the boolean AND operator following the formula: CD4 AND CD40L-CD4 AND [cytokine]-CD4, applied separately for IL-2, TNF-α, MIP-1β, and IFN-γ. CD40L was poorly expressed on $CD8^+$ T cells and was therefore not assessed in this subset. Accordingly, individual cytokine/CD8 gates defined each cytokine-producing $CD8^+$ T-cell population. Cytokine co-expression was not analyzed owing to the absence of detectable HIV-specific T-cell responses.

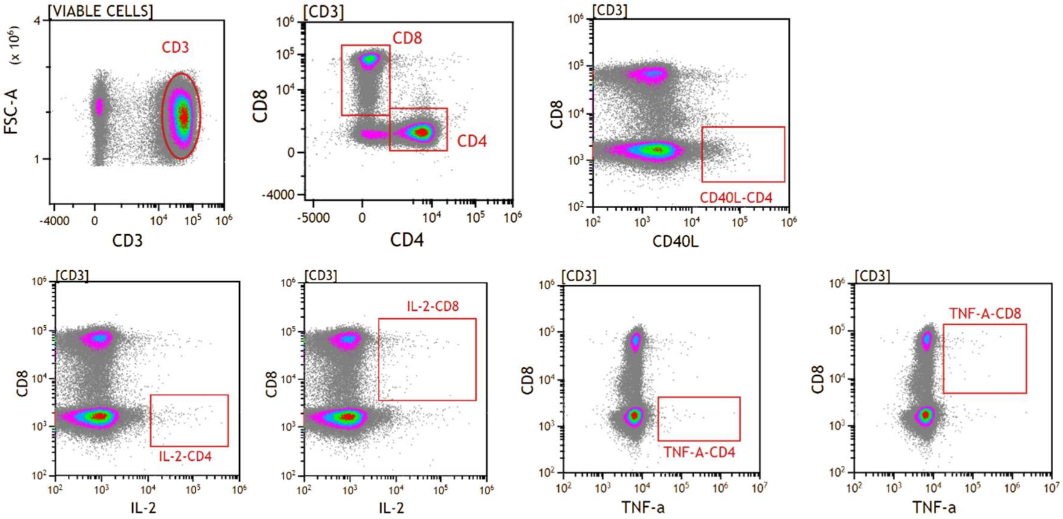

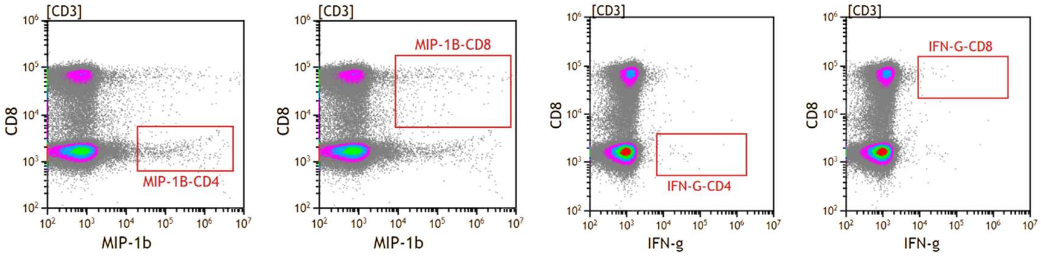

### Supplemental information – Part 2: Data

#### Table S2-1. Characteristics of 76 participants included in the ANRS-EP59-CLEAC study.

| Characteristics [a] | Median [IQR]<br>% (n) |
|---|---|
| Age, years | 11 [8;14] |
| Sex | |
| Male | 46.0 (35) |
| Female | 54.0 (41) |
| Sub-Saharan African origin | |
| No | 25.0 (19) |
| Yes | 75.0 (57) |
| Born in mainland France | |
| No | 40.8 (31) |
| Yes | 59.2 (45) |
| Age at ART1 initiation (months) | 25.2 [2.3 ;79.0] |
| Time since ART1 initiation (months) | 93 [57;137] |
| Current HIV RNA | |
| < 50 copies/mL | 82.9 (63) |
| ≥ 50 copies/mL | 17.1 (13) |
| CD4 count, cells/µL | 856 [685;1236] |
| CD4 percentage | 38 [32.5;42] |
| CD8 count, cells/µL | 714 [519.5;985] |
| CD8 percentage | 29 [25;36] |
| CD4/CD8 ratio | 1.34 [0.92;1.76] |

[a] adapted from [8].

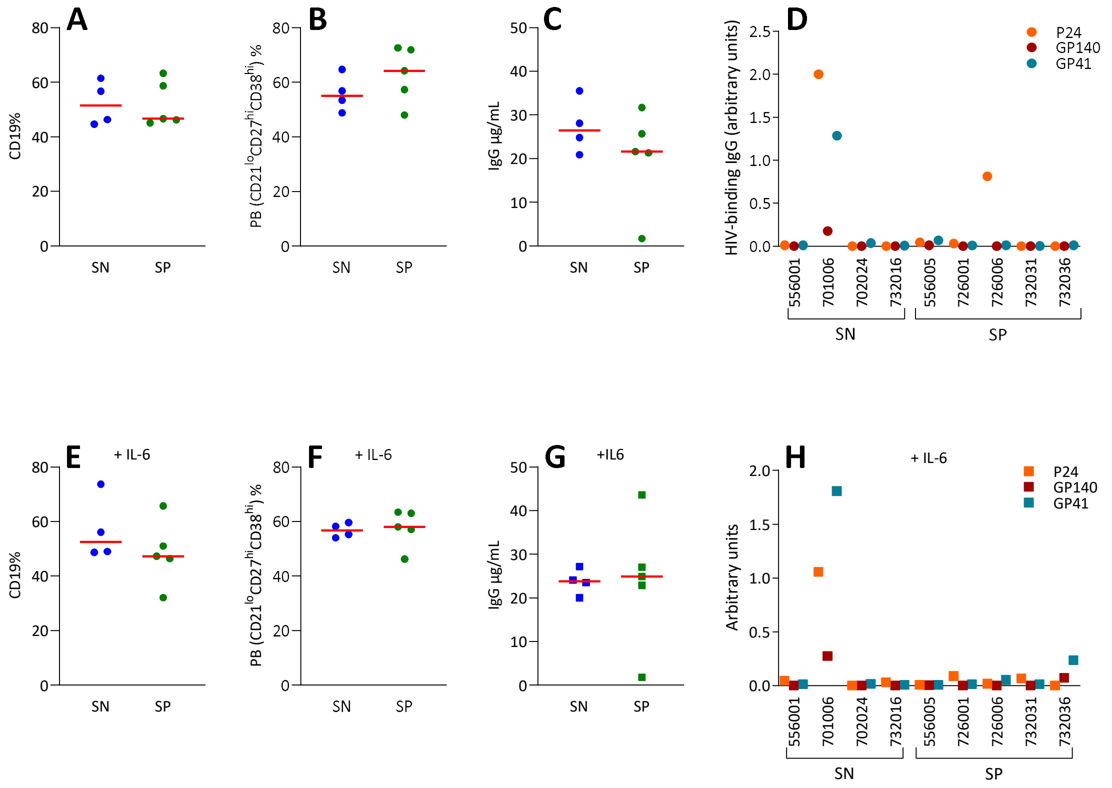

#### Figure S2-1. IL-6 supplementation did not enhance memory B cell stimulation or HIV-specific antibody production *in vitro*.

We thawed PBMCs, stimulated them with the TLR7/8 agonist R848, and cultured them with IL-2 (panels A to D) or IL-2 + IL-6 (panels E to H). For comparison, we reproduced data from IL-2-supplemented cultures (panels A to D) from Figure 3. Flow cytometry after 7 days of culture determined the percentages of B lymphocytes (CD19+, panels A and E) among viable cells and plasmablasts ($CD19^{+}CD21^{lo}CD27^{hi}CD38^{hi}$, panels B and F) among B cells. At day 12, we collected supernatants to quantify total IgG (panels C and G) and HIV-specific IgG by ELISA (panels D and H). Red bars indicate medians and *P* values above graphs correspond to Mann-Whitney tests.

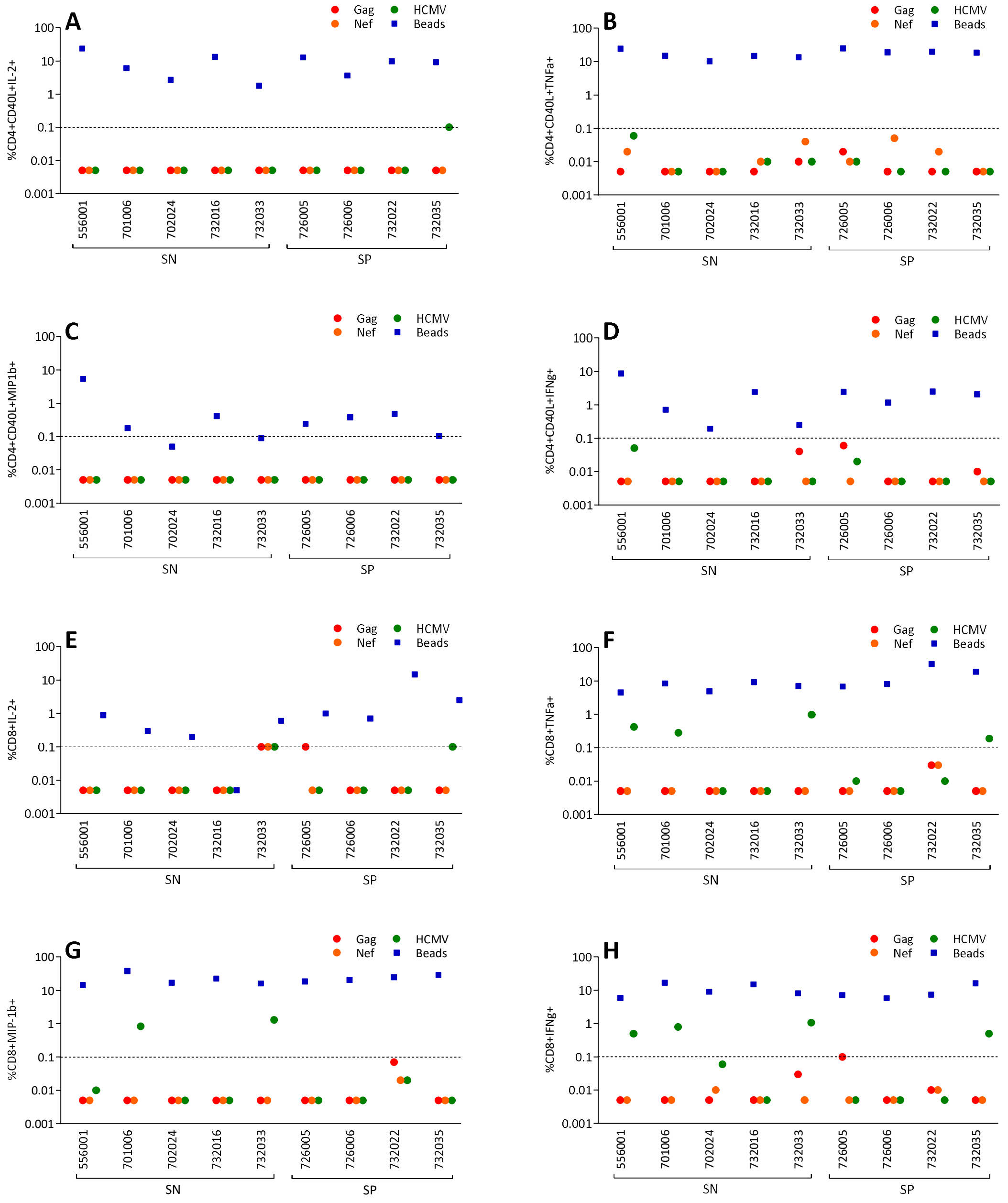

#### Figure S2-2. HIV-specific T cells were undetectable by intracellular flow cytometry assay.

We thawed frozen PBMCs and stimulated them with peptide pools or beads coated with anti-CD3 and anti-CD28 antibodies in the presence of brefeldin A. CD3, CD4, CD8, CD40L, IL-2, TNF-α, MIP-1β and IFN-γ molecules were quantified by flow cytometry. Panels A to D present the frequencies of CD4 T lymphocytes coexpressing CD40L and each of the four cytokines. Panels E to H present the frequencies of CD8 T lymphocytes expressing each of the four cytokines. Dotted lines indicate the positivity threshold. Data are shown for nine children, 5 SN and 4 SP.

Figure S2-3. NK lymphocyte subsets did not associate with negative HIV serology

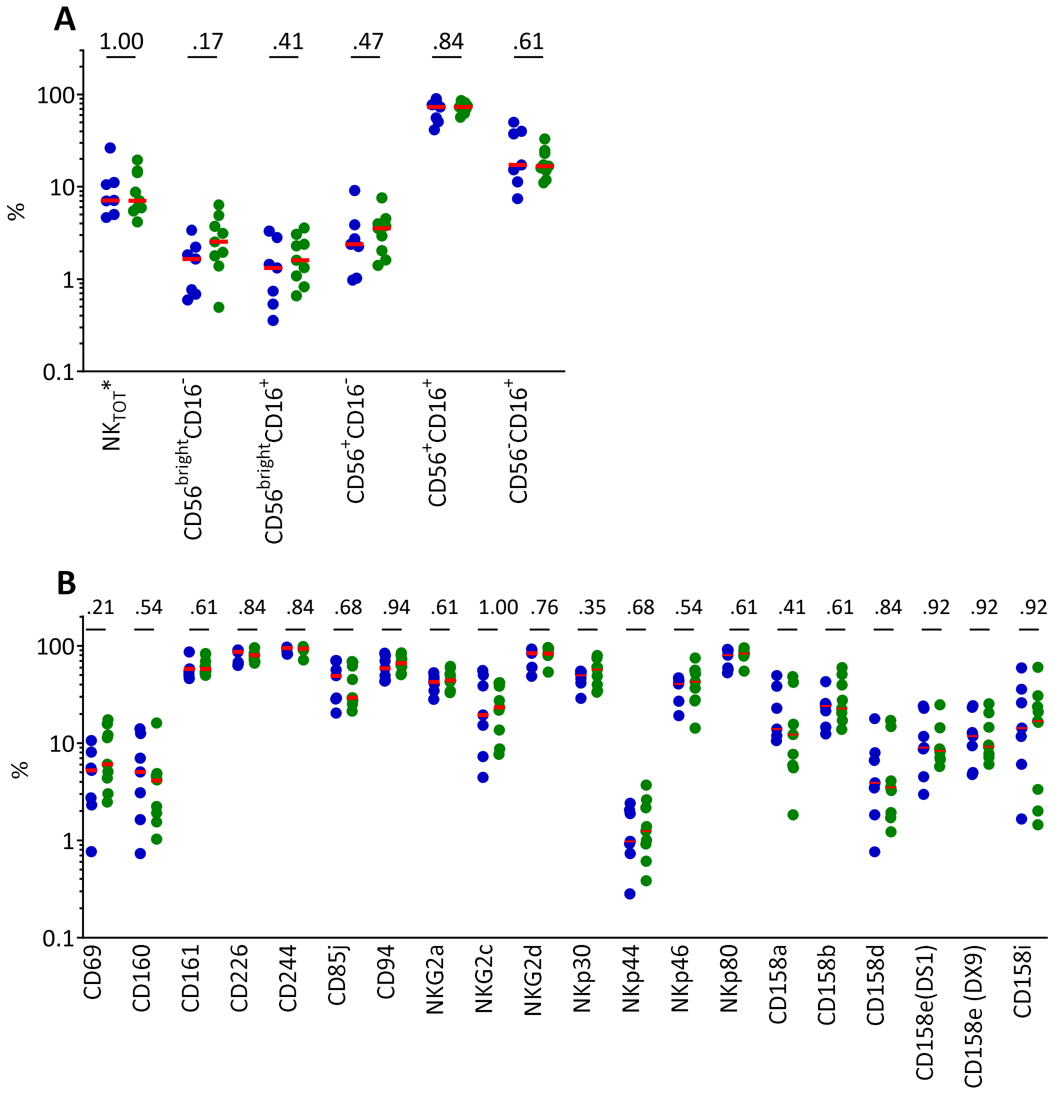

**Supp Figure 3**

Figure S2-3. NK lymphocyte subsets did not associate with negative HIV serology. We analyzed fresh blood cells by flow cytometry to quantify NK lymphocytes. Among early-treated children with sustained HIV suppression, we compared NK cells between SN and SP. A. NK cell levels (% among lymphocytes) and five subsets defined by CD16 and CD56 expression (% among NK cells). B. The CD69 activation molecule and stimulatory and inhibitory NK receptors (% of cells expressing markers among total NK cells). Red bars indicate medians and *P* values above graphs correspond to Mann-Whitney tests.

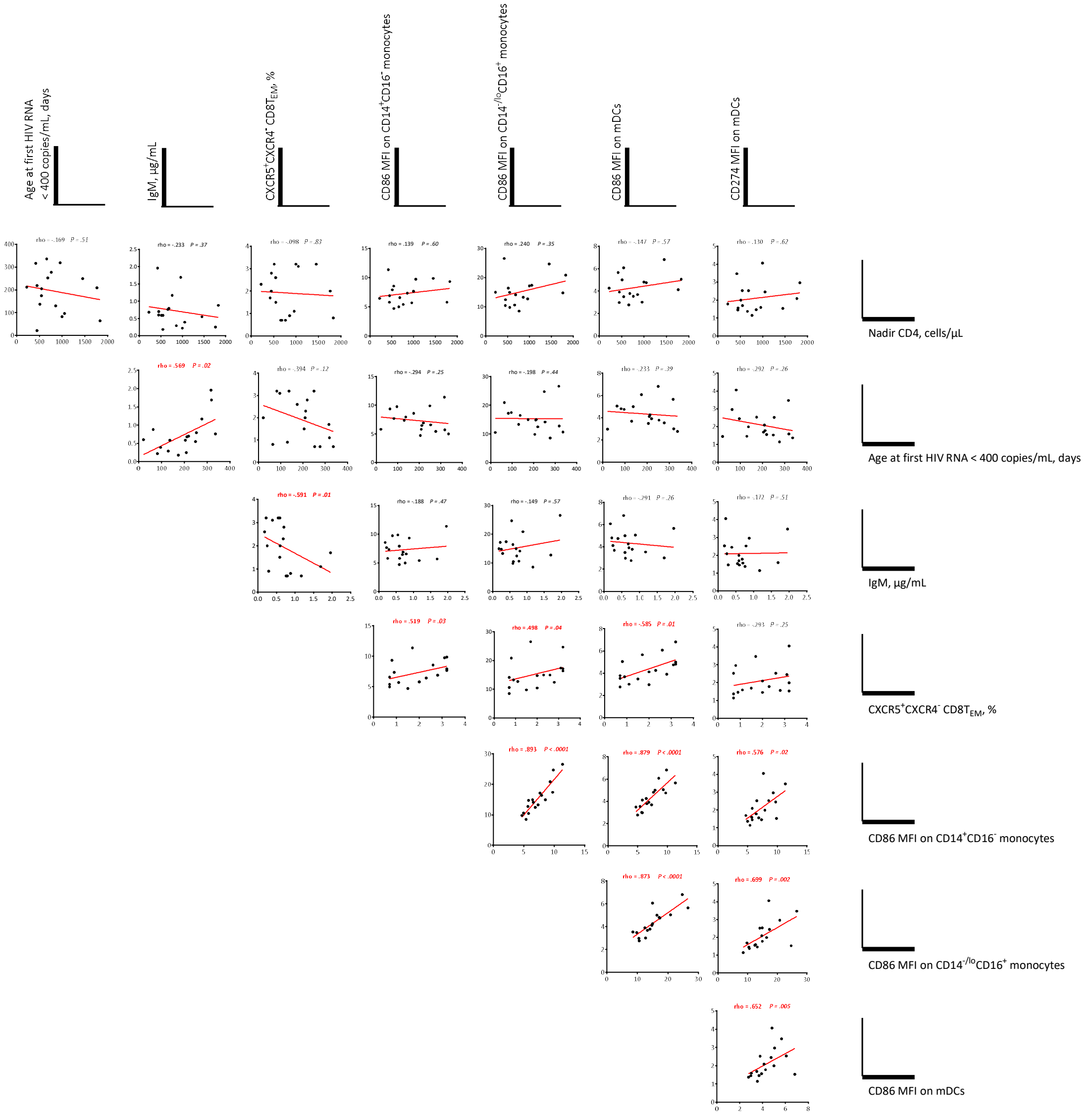

#### Figure S2-4. Correlations between variables associated with negative HIV serology.

We used the Spearman test to assess correlations between continuous variables which were associated with negative HIV serology (Age VS, Age at first HIV RNA < 400 copies/mL, days; IgM, µg/mL; $X5^{+}R4^{-}$ $CD8T_{EM}$, $CXCR5^{+}CCR4^{-}$ $CD8T_{EM}$, %; $CD86\_CD14^{+}16^{-}M$, CD86 MFI on $CD14^{+}CD16^{-}$ monocytes; $CD86\_CD14^{-}16^{+}M$, CD86 MFI on $CD14^{-/lo}CD16^{+}$ monocytes; CD86_mDC, CD86 MFI on mDCs ; CD274_pDC, CD274 MFI on pDCs; nadCD4, Nadir CD4, cells/µL). Coefficients and *P* values are indicated above graphs. Bold characters indicate *P* values < 0.05.
